# Evolutionarily conserved transcriptional regulators control monoaminergic neuron development

**DOI:** 10.1101/2025.10.29.685200

**Authors:** Clifton Lewis, Matthew Goulty, Aniela Wroblewska, Nicola Croxall, David Onion, Sue Robinson, Robert P. Zinzen, Jordi Solana, Charlambos P. Kyriacou, Ezio Rosato, Roberto Feuda

**Affiliations:** Neurogenetics Group, Division of Genetics and Genome Biology, School of Biology and Biomedical Sciences, College of Life Sciences, University of Leicester, Leicester LE1 7RH, UK; Centre for Chromosome Biology, Biomedical Sciences Building, University of Galway, Galway, H91 W2TY, Ireland; The Francis Crick Institute, London, UK; Faculty of Medicine & Health Sciences, Queen’s Medical Centre, Nottingham NG7 2UH, UK; Division of Molecular and Cell Biology, College of Life Sciences, University of Leicester, Leicester LE1 7RH, UK; Systems Biology Imaging Technology Platform, Max Delbrück Centre for Molecular Medicine (MDC-BIMSB), Berlin, Germany; Living Systems Institute, University of Exeter, Exeter, UK; Department of Biosciences, University of Exeter, Exeter, UK; Department of Biology, Geology and Environmental Science, University of Bologna, Bologna, Italy

## Abstract

To what extent conserved developmental programs specify homologous cell types is a central question in biology. Here, we address this by focusing on reconstructing monoaminergic neuron development in *Drosophila melanogaster* embryo using time- resolved single-cell genomics, spatial transcript mapping with hybridisation chain reaction, and targeted metabolomics. We uncover a regulatory landscape in which specific transcription factors are activated before biosynthetic enzymes, establishing a prospective temporal architecture for monoaminergic fate specification. Comparative analyses of developmental single-cell atlases from zebrafish and sea urchin indicate that components of this machinery are conserved across ∼550 million years of bilaterian evolution with orthologous transcription factors showing similar temporal dynamics. Together, these findings point to a putatively conserved regulatory core that interfaces with other context-dependent transcription factors; this interplay accommodates monoaminergic multifunction and subtype diversity across distinct neuroanatomies.

## Introduction

Understanding whether conserved or divergent developmental programs underlie the specification of homologous cell types is a central goal in evolutionary and developmental biology. We address this question by focusing on monoaminergic neurons, a class of cells that produce neuromodulators such as dopamine and serotonin. Monoaminergic neurons provide a powerful system to study evolution and development for several reasons. They are molecularly defined by the expression of key biosynthetic enzymes (Libersat and Pflueger, 2004), they are widespread across bilaterians, their monoamine products can be precisely quantified, and the genes underlying their biosynthetic pathways have been shown to derive from a common evolutionary origin (Parent, 1984; Yamamoto and Vernier, 2011; Goulty *et al*., 2023).

Although these neurons are extremely rare (<1% of neurons in fly and mammalian brains), they play fundamental roles in processes ranging from motor control and circadian rhythms to learning and mood (Libersat and Pflueger, 2004; Flames and Hobert, 2011; Monti, 2011; Waddell, 2013; Zheng *et al*., 2025). Furthermore, their dysfunction contributes to major human neurological and psychiatric disorders, including Parkinson’s disease, depression, and schizophrenia (Kurian *et al*., 2011; Volkow *et al*., 2011; Grace, 2016).

The developmental specification of monoaminergic neurons requires the coordinated activity of multiple transcription factors (TFs) and other regulatory genes organised into hierarchical gene regulatory networks (GRNs) (Hobert, 2008; Flames and Hobert, 2009, 2011). Studies across vertebrates and invertebrates have identified TFs that are critical for establishing monoaminergic fate (Flames and Hobert, 2011). In vertebrates, for example, LIM-homeodomain proteins Lmx1a/b, the ETS factor Fev/Pet-1, and Nkx2.2 are essential for serotonergic neuron specification (Cheng *et al*., 2003; Ding *et al*., 2003; Hendricks *et al*., 2003). In *Drosophila*, the zinc-finger proteins Eagle (eg) and Huckebein (hkb) play similar roles in serotonergic differentiation (Dittrich *et al*., 1997). In *C. elegans*, dopaminergic fate is controlled by terminal selector TFs such as the ETS transcription factor, AST-1 (Flames and Hobert, 2009). Previous work from our lab established that monoaminergic cell identity is defined by a conserved set of transcription factors shared across bilaterian species, suggesting that a common regulatory basis for these neurons originated early in animal evolution (Goulty *et al*., 2025). Yet these candidate-based studies provide only a partial view of the regulatory landscape underlying monoaminergic specification and their possible conservation. We therefore still lack a systems-level understanding of how monoaminergic neurons develop. Addressing these questions is essential for understanding whether homologous cell-types develop following a conserved transcriptional program.

Here to address this open question, we combine time-resolved single-cell transcriptomics, trajectory modelling, spatial transcript validation, and targeted metabolomics in embryonic neurogenesis of *Drosophila melanogaster*, and test for evolutionary conservation in zebrafish and sea urchin development. Our results reveal the developmental program of monoaminergic neurons in the *Drosophila* embryo, including key TFs—*fd59A*, *dmrt99B*, and *Lmx1a*—that pattern these lineages and differences in maturation rates between subtypes. We also identified several TF orthogroups conserved across *Drosophila*, sea urchin, and zebrafish. Comparison of monoaminergic developmental trajectory between *Drosophila* and zebrafish revealed a similar developmental cascade of TFs. Together, these findings support a putative conserved temporal logic for monoaminergic fate acquisition and begin to explain how developmental fate and function can align and/or diverge across species.

## Results

### A temporally resolved single-cell atlas captures neurogenesis in *Drosophila*

To understand the development of monoaminergic neurons, we performed single-cell RNA sequencing (scRNA-seq) on *Drosophila melanogaster* embryos collected at eight partially overlapping time windows spanning 0 to 22 hours after egg laying (AEL), with two biological replicates per window (**Figure 1A**.), covering the transition from neuroectoderm to fully differentiated embryonic neurons. To enrich for progenitor and early neural population, we used a SoxN-sfGFP transgenic line (VDRC #318062, Sarov *et al*., 2016) to isolate proneural cells and neuroblasts, and anti-Elav immunostaining to enrich for differentiating neurons, followed by fluorescence-activated cell sorting (FACS) (**Figure S1.**). Single cell RNA-seq libraries were prepared on the 10X Genomics v3.1 platform and sequenced using Illumina NovaSeq X Plus.

**Figure 1.**
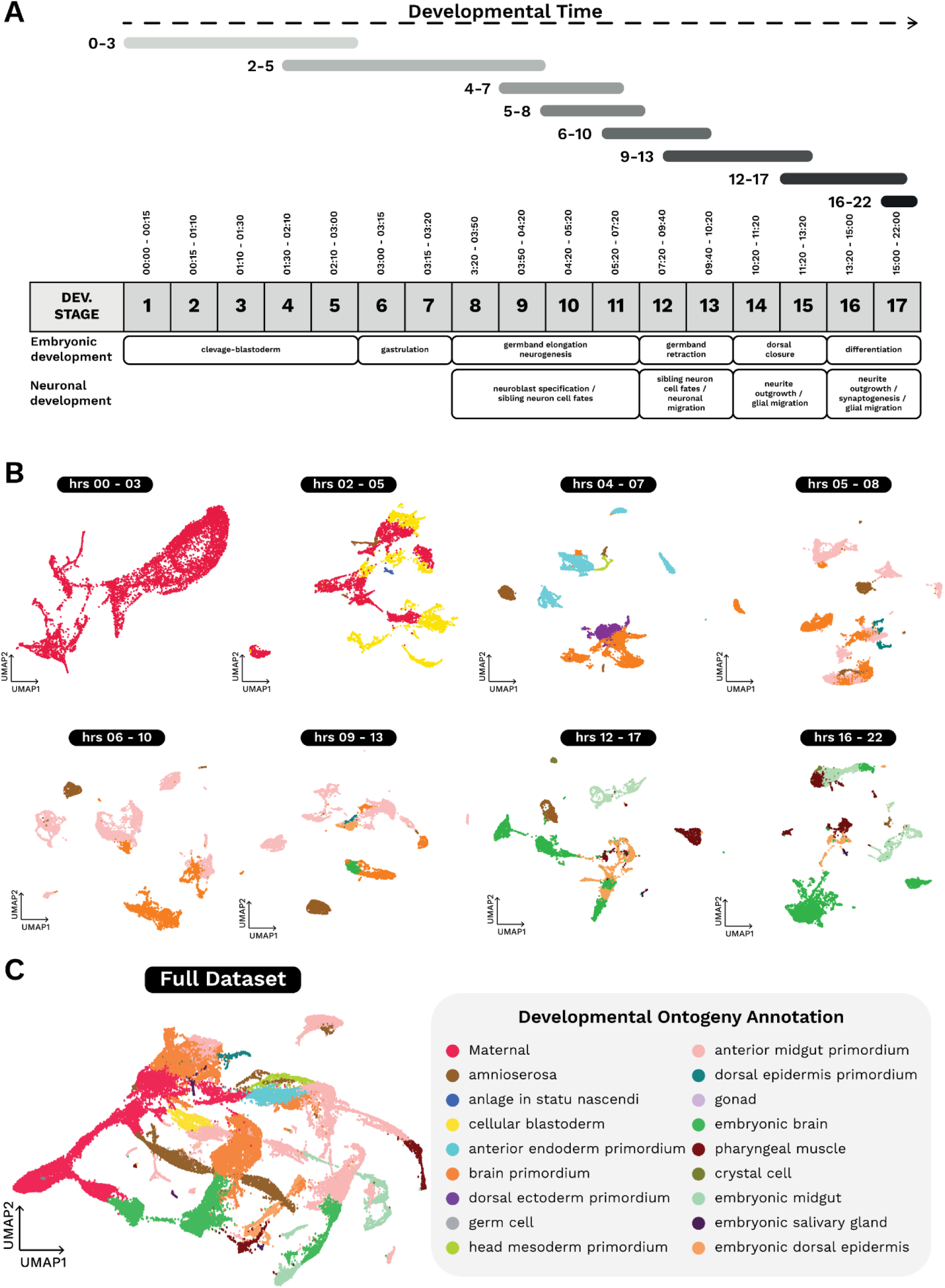
Time-resolved sampling strategy and developmental ontogeny annotation of targeted scRNA-seq across *Drosophila* embryogenesis. **A.** Timeline of eight partially overlapping collection windows, spanning embryogenesis, from 0–22 hrs after egg laying (AEL) aligned to canonical embryonic stages (Bownes stages 1–17, (Campos-Ortega and Hartenstein, 1985)) and key morphogenetic landmarks (cleavage/blastoderm, gastrulation, germband elongation, germband retraction, dorsal closure, and differentiation). Major neurogenic milestones are indicated below the timeline, including neuroblast specification, diversification of sibling neuron fates, neuronal migration, neurite outgrowth, glial migration, and synaptogenesis. **B.** UMAPs for each time window showing single-cell transcriptomes coloured by developmental ontology. **C.** UMAP of the combined dataset with cells coloured by developmental ontology derived from *in situ*–guided annotation. Representative labels include maternal, cellular blastoderm, amnioserosa, dorsal ectoderm primordium, brain primordium, embryonic brain, and embryonic midgut.

After standard quality control, we retained 57,476 high-quality cells, resolving into transcriptionally distinct clusters. Cluster identities were assigned by mapping cluster-specific marker genes—selected as the top differentially expressed features with adjusted p-value < 0.05—to *in situ* expression patterns from the Berkeley Drosophila Genome Project (BDGP) using an annotation pipeline adapted from Calderon *et al*. (2022). This analysis revealed a clear temporal transition: early clusters were enriched for maternal and early zygotic genes, whereas later stages featured organised, differentiated tissues; including the brain primordium, embryonic brain, and embryonic gut (**Figure 1B**. and **Table S1.**). Embedding all time points together revealed a continuous developmental trajectory (**Figure 1C**.), capturing the progressive emergence of neuronal identity.

### Neurotransmitter subtype emergence reveals early co-expression and lineage complexity

Although our dataset is enriched for neural progenitors and differentiated neurons, cluster annotations revealed the presence of additional cell types (**Figure 1C**.). This likely reflects autofluorescence and/or putatively *SoxN*-driven GFP expression in the developing gut, leading to non-specific capture during FACS enrichment (**Figure S1.**).

To resolve distinct stages of neurogenesis, we used established markers - e.g. *dpn*, *wor*, *mir*, *elav*, *nSyb* - to identify neuroblasts, ganglion mother cells (GMCs), intermediate neural progenitors (INPs), and mature neurons in a population of ∼40,000 cells (**Figure 2A**., **Figure S3A.**) (Dillon *et al*., 2022; Peng *et al*., 2024). To capture the full neurogenic continuum, spanning from neuroblasts to mature neurons, we subsampled to ∼10,000 high-quality cells, excluding disparate clusters (**Figure 2B**., **Figure S3B.**). These cells were distributed across all the developmental time points, enabling the reconstruction of continuous differentiation trajectories (**Figure 2C**.). Among these terminal-state mature neurons, we identified seven neurotransmitter-defined populations: cholinergic (931 cells), GABAergic (816 cells), glutamatergic (173 cells), monoaminergic (77 cells), and three mixed populations co-expressing markers of GABA and glutamate (100 cells), acetylcholine and GABA (556 cells), or serotonin and GABA (388 cells), respectively (**Figure 2B**.).

**Figure 2.**
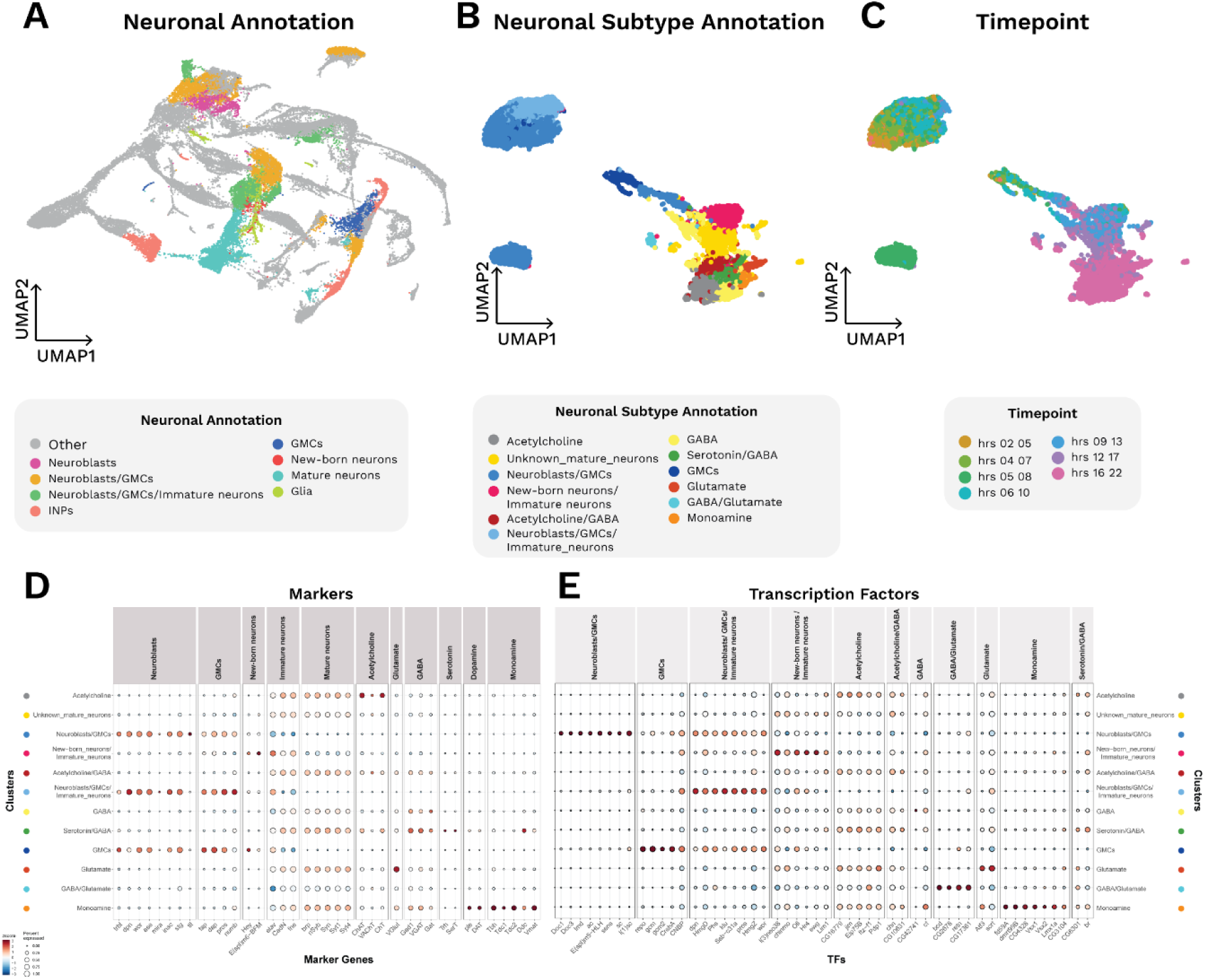
Time-resolved single-cell landscape of embryonic neurogenesis in ***Drosophila*** **A. Neuronal annotation.** UMAP of the full embryonic dataset with neural lineages highlighted and non-neuronal cells in grey. Colours denote major neural states: neuroblasts, neuroblasts/GMCs, neuroblasts/GMCs/immature neurons, intermediate neural progenitors (INPs), GMCs, new-born/immature neurons, mature neurons, and glia. **B. Neuronal subtype annotation.** Subclustered neurogenic atlas reannotated to subtypes including transmitter-class/subtype as well as neuronal developmental intermediates. Classes include acetylcholine, GABA, glutamate, monoamine, mixed identities (e.g., acetylcholine/GABA; GABA/glutamate), unknown mature neurons, and progenitor states (neuroblasts/GMCs; neuroblasts/GMCs/immature neurons; GMCs; new-born/immature neurons). **C. Timepoint.** Subclustered dataset coloured by collection window, showing temporal separation across the UMAP. **D. Markers.** Dot plot of selected marker genes across the neuronal states/subtypes in the subclustered dataset. Dot size indicates the fraction of cells expressing the gene; colour reflects average expression. **E. Differential transcription factors.** Dot plot of differential TFs across neuronal states/subtypes, highlighting regulators associated with progenitors, early differentiation, transmitter specification, and mature neurons.

Given that distinct TF combinations are known to contribute to neuronal subtype identity, we next sought to identify TFs associated with each neurotransmitter-defined population (**Figure 2E**.). Our differential TF expression analysis revealed that, with the exception of neuroblasts/GMCs, GABA/glutamatergic neurons, and monoaminergic neurons, most of these neuronal populations could not be unambiguously resolved based on TF expression alone. This outcome suggests that TFs in these other lineages may lack strict subtype specificity, that neuronal identity may arise from more complex regulatory interactions beyond TF expression, or that the dataset may not have sufficient depth to fully resolve these distinctions. Monoaminergic neurons, however, showed co-expression of multiple TFs, including *fd59A*, *dmrt99B*, *Lmx1a*, and *Vsx1* and *Vsx2* (**Figure 2E**.). In contrast, the mixed serotonergic/GABAergic population was marked by the co-expression of *CG8301* and *br* (**Figure 2E**.). These findings define candidate regulatory genes underlying monoaminergic fate.

### RNA velocity reveals continuous differentiation trajectories

To investigate the transcriptional dynamics underlying monoaminergic neuron differentiation, we applied RNA velocity analysis using scVelo (La Manno *et al*., 2018; Bergen *et al*., 2020) and modelled lineage trajectories with CellRank (v2) (Lange *et al*., 2022; Weiler *et al*., 2024), which integrates splicing kinetics (Velocity Kernel), transcriptomic similarity (Connectivity Kernel), and CytoTRACE dynamics to infer directed developmental transitions (CytoTRACE Kernel). We first applied scVelo to the full dataset. Velocity vector fields projected onto the UMAP embedding revealed a strong correlation between transcriptional dynamics and the temporal structure of the dataset (**Figure S4.**).

We then focused on the subclustered population spanning the full neurogenic trajectory from neuroblasts to mature neurons (**Figure 2B–D**.). Application of scVelo and CellRank to this dataset recovered clear trajectories for each subtype and a smooth latent-time ordering (**Figure S5A.**). To interpret the underlying programs, we identified velocity-inferred dynamical genes with the scVelo dynamical model and ordered cells by latent time (**Figure S5B.**). We discretised latent time into 20 bins and, for each bin, compiled the genes whose fitted expression/velocity profiles peaked within that interval. Gene Ontology (GO) over-representation analyses (ORA) were then performed using all expressed genes in the subset as background with Benjamini–Hochberg correction. The enrichment landscape shows a coherent progression: (i) early bins are dominated by cell-state remodelling and tissue-building programs—RNA processing, cell cycle/proliferation, morphogenesis, and organogenesis; (ii) intermediate bins transition toward neuronal differentiation, with terms related to synapse formation and cell morphogenesis involved in neuron differentiation; while (iii) late bins are enriched for synaptic and excitability modules, including neuron recognition, axon guidance, and neuroactive ligand–receptor interaction, consistent with the acquisition of functional neuronal properties. Together, these GO trends align with the velocity-derived latent-time ordering and provide functional support for a continuous maturation from progenitor states to functionally specialised neurons (**Figure S5C.**).

To reconstruct the differentiation trajectory of monoaminergic neurons in greater detail, we applied CellRank to define initial and terminal states and to model transcriptional dynamics within this specific lineage (**Figure 3A**.). Genes encoding key enzymes required for monoaminergic neurotransmission—including *ple*, *Tdc1*, *Tdc2*, *Tbh*, and *Vmat*—were strongly upregulated toward the terminal state, as expected, marking the final stages of differentiation. With the exception of *Trhn* and *Ddc*, these enzymes exhibited high specificity for the monoaminergic lineage (**Figure S6.**), indicating subtype-restricted induction of the biosynthetic program. Similarly, top-ranking TFs associated with monoaminergic identity—including *fd59A*, *dmrt99B*, and *Lmx1a*—were also selectively expressed in the monoaminergic trajectory (**Figure 3B**.) and peaked at approximately the same latent time as the biosynthetic genes (**Figure 3A**.). This coordinated expression suggests a tightly coupled regulatory program in which TFs may directly activate terminal differentiation effectors. In contrast—and consistent with their role as responders to monoamine signals—genes encoding monoaminergic receptors showed broader expression across multiple neuronal lineages (**Figure S7.**). This pattern suggests that receptor deployment is decoupled from biosynthetic identity and instead reflects widespread postsynaptic responsiveness to monoamines.

**Figure 3.**
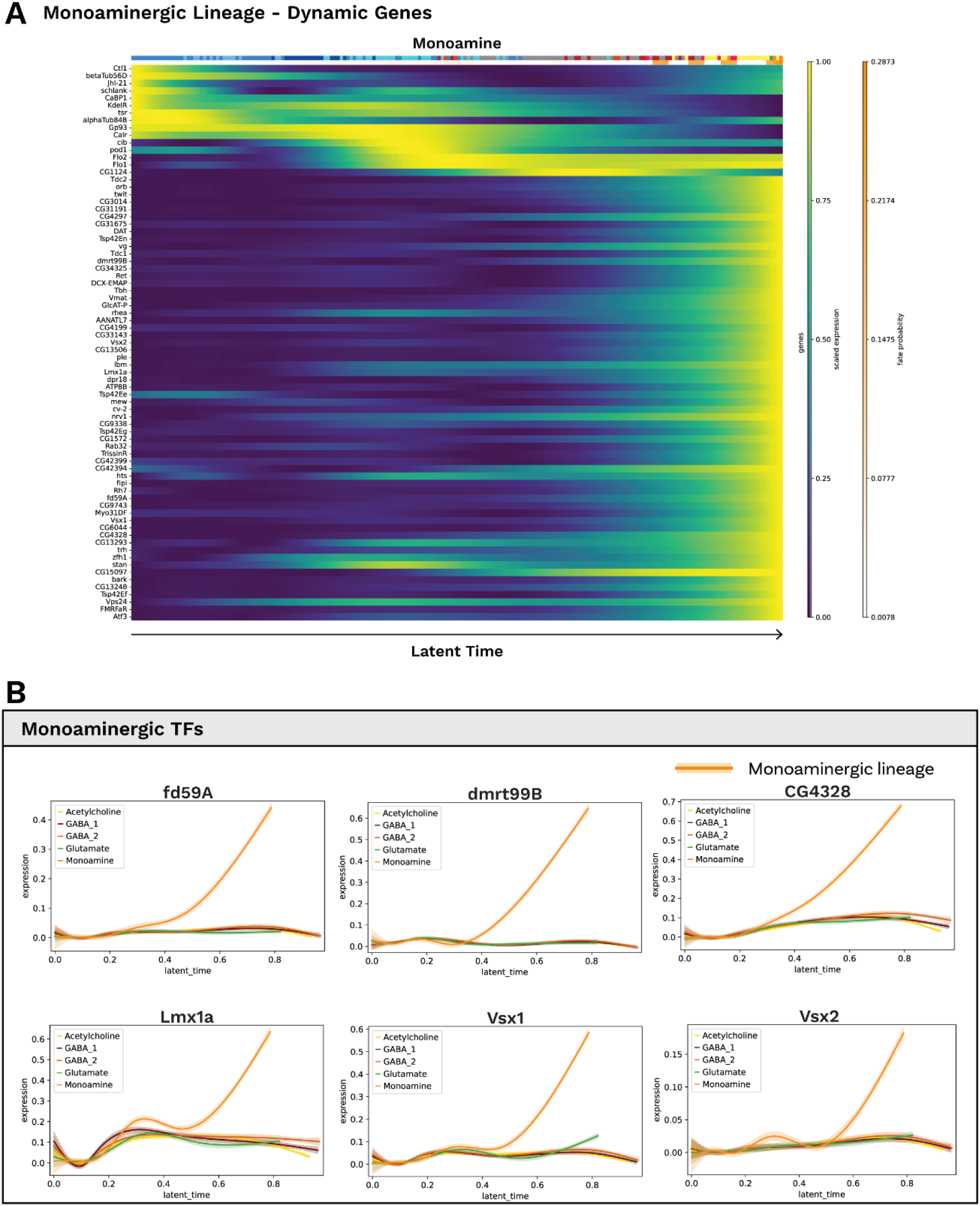
Monoaminergic lineage—dynamic gene programs and candidate transcription factors gene trends **A. Dynamic genes along latent time.** Heatmap of CellRank-identified dynamical genes within the monoaminergic lineage. Cells are ordered by latent time (left→right; arrow) and expression is z-scaled per gene (yellow = higher expression). Many neurotransmitter-pathway genes and regulators turn on progressively toward late latent time as well as many of the candidate differentially expressed TFs identified previously. **B. Monoaminergic transcription factors.** Smoothed expression trajectories across latent time for selected TFs (*fd59A*, *dmrt99B*, *CG4328*, *Lmx1a*, *Vsx1*, *Vsx2*). The monoaminergic lineage is highlighted in orange; other neuronal lineages (acetylcholine, GABA_1/2, glutamate) are shown for comparison. Curves show lineage-wise mean trends (y-axis, scaled expression) across latent time (x-axis), illustrating the preferential late-stage upregulation of these TFs in the monoaminergic lineage specifically.

### Spatial validation supports coordinated transcriptional activation and resolves monoaminergic subtype identity

Although our scRNA-seq atlas resolves the major neuronal lineages and transmitter classes, monoaminergic neurons are a rare population in the *Drosophila* CNS (Busch *et al*., 2009; Hartenstein *et al*., 2017; Kasture *et al*., 2018; Babski, Codianni and Bhandawat, 2024). As a result, it was difficult to clearly separate specific monoaminergic subtypes from transcriptomes alone. To both validate CellRank-inferred dynamics and add spatial specificity, we focused on two subtypes – serotonergic and dopaminergic - and performed hybridisation chain reaction (HCR) (Choi *et al*., 2018).

We assayed a panel that included most of the monoaminergic biosynthetic enzymes, their receptors, and six transcription factors (**Figure 4**., **Figure S8-9.**). TFs were chosen based on differential expression within the monoaminergic lineage (**Figure 2C**.) and prior evidence for roles in monoaminergic fate. Spatial profiling showed that *fd59A*, *dmrt99B*, and *Lmx1a* are expressed in the CNS beginning at stages 10–12 (∼8 h AEL), preceding the onset of biosynthetic gene expression. In contrast, monoaminergic enzymes were robustly expressed from stage 12 (∼10 h AEL) onward, consistent with the timing of neuronal specification. Notably, this TF induction occurs slightly earlier than suggested by our latent-time analysis, which predicted a more coordinated co-activation with the enzymes. Receptors generally followed a similar onset, with most appearing from stage 12. A notable exception was *5-HT2A*, which was detected as early as stage 1 and in non-neural tissues, consistent with a recent study implicating it in early morphogenetic roles beyond neurotransmission (Karki *et al*., 2023).

**Figure 4.**
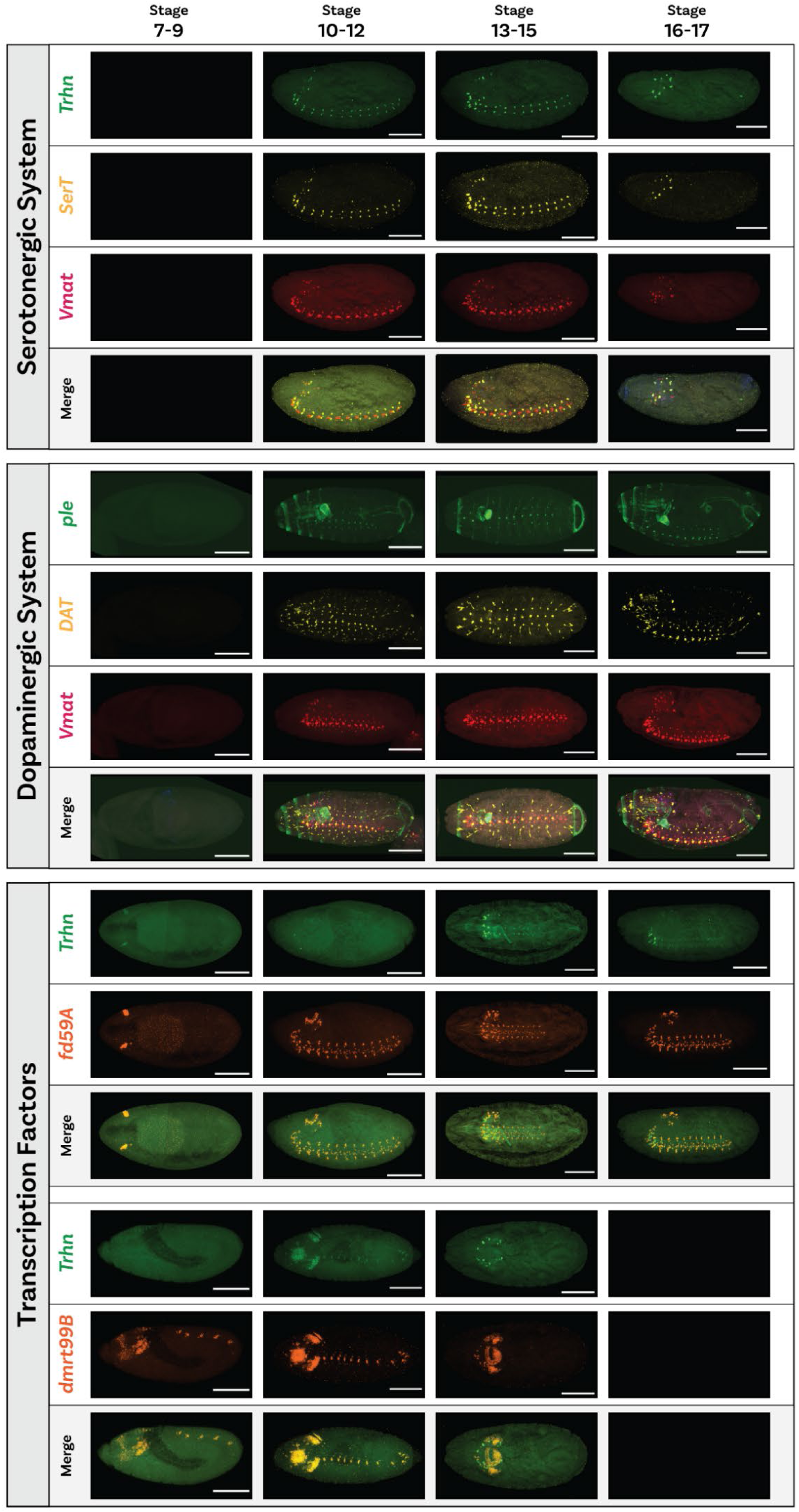
HCR validates the temporal onset and spatial organisation of serotonergic and dopaminergic programs in the embryo HCR staining across four developmental windows (columns: stages 7–9, 10–12, 13– 15, 16–17). Each image shows a whole-mount embryo, lateral view with anterior to the left. Scale bars 100 μm. **Serotonergic system**. HCR for *Trhn* (*tryptophan hydroxylase*, green), *SerT* (*serotonin transporter*, yellow) and *Vmat* (*vesicular monoamine transporter*, magenta), plus merges. No signal is detected at stages 7–9. From stage 10–12 onwards *Trhn*, *SerT* and *Vmat* appear in segmentally repeated CNS domains; marking serotonergic neurons as they mature. **Dopaminergic system**. HCR for *ple* (*Tyrosine hydroxylase*, green), *DAT* (*dopamine transporter*, yellow) and *Vmat* (magenta), plus merges. *ple* is first detected by stage 10–12, with *DAT* and *Vmat* also appearing by stage 10-12. *ple*/*DAT* signals co-localise with *Vmat* in the CNS. **Transcription factors**. HCR for *fd59A* or *dmrt99B* (orange) together with *Trhn* (green), plus merges. TF expression is evident in the CNS earlier (stages 7-9) and broadens by stages 10-12, preceding with enzyme onset, consistent with a putative upstream regulatory role in monoaminergic fate.

Together, these data mostly corroborate the temporal ordering inferred by CellRank and, crucially, resolve subtype identity by localising expression to serotonergic or dopaminergic domains. They further suggest that these TFs, *fd59A*, *dmrt99B*, and *Lmx1a*, may act upstream to specifically initiate part of the monoaminergic program, with effector enzymes and receptors switching on subsequently.

### Mass spectrometry reveals temporally distinct onset of monoamine production

Our transcriptional and spatial analyses suggests that by stage 12 (∼9–10 h AEL), the *Drosophila* embryo expresses the biosynthetic enzymes and receptors required to produce and respond to monoamines. However, whether these neurons are functionally active at this stage remains unresolved. Previous studies have detected serotonin and dopamine via immunohistochemistry only from stage 17 (∼15 h AEL) onwards (Lundell and Hirsh, 1994), raising questions about the relationship between transcriptional commitment and neurotransmitter availability. To resolve this dichotomy, we conducted targeted mass spectrometry across seven developmental windows spanning 0–22 h AEL. At each time point, we analysed ≥6 biological replicates (54 samples total) and quantified 19 analytes, including all canonical monoamines (**Table S2.**) GABA, glutamate, and acetylcholine **(Figure S11.).**

Our results revealed distinct temporal profiles across monoaminergic subtypes (**Figure 5.; Figure S10.**). Histamine and tyramine were constitutively present throughout development, suggesting their production may occur independently of canonical neurogenesis. In contrast, both dopamine and serotonin exhibited temporally restricted accumulation, with sharp increases after stage 12—closely aligning with the transcriptional induction of their biosynthetic enzymes. Although dopamine and serotonin followed similar temporal trajectories, their relative levels differed, likely reflecting variations in biosynthetic efficiency or substrate availability. It is important to note that this effect may also reflect differences in the detection capabilities for these molecules.

**Figure 5.**
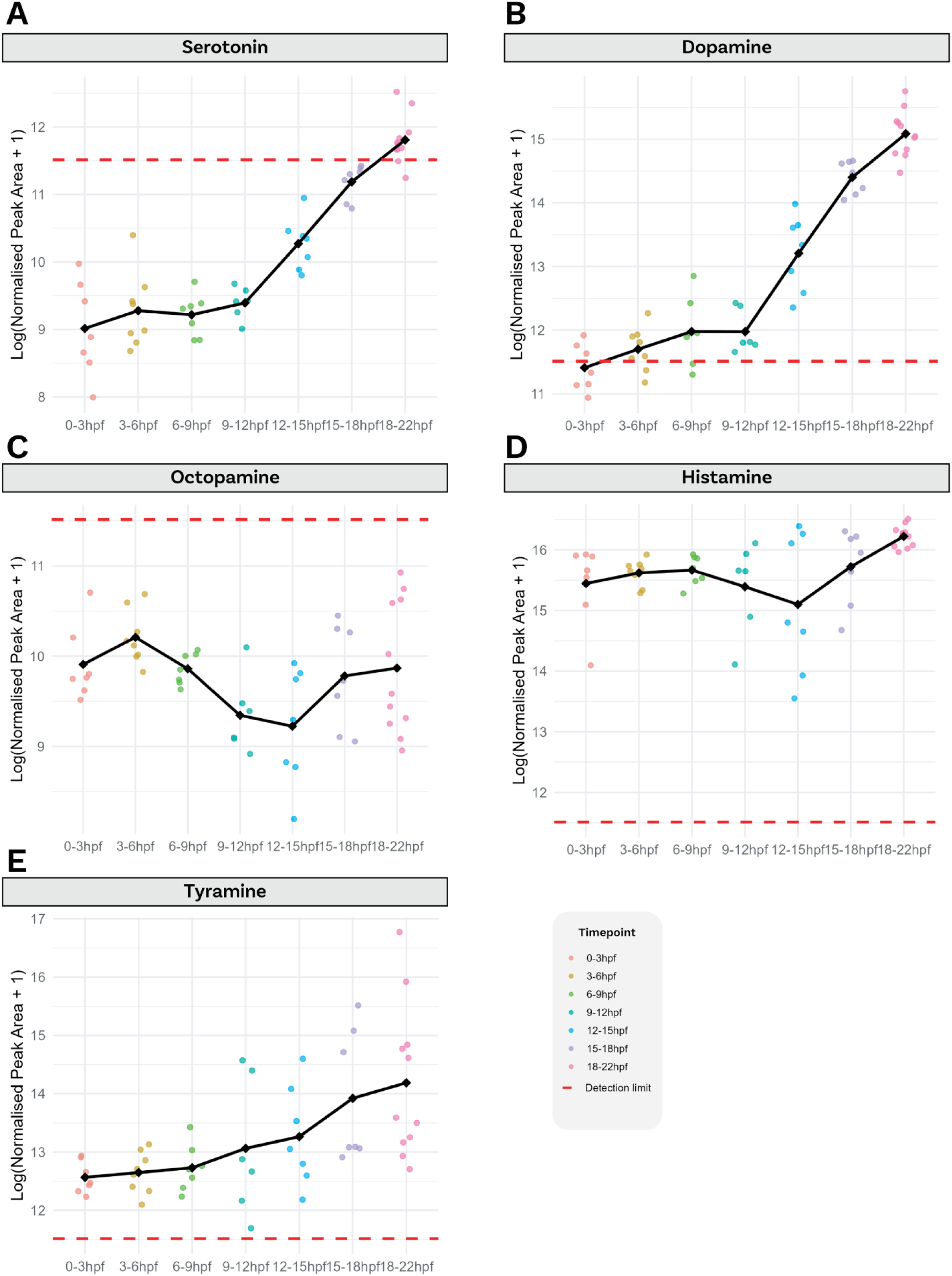
Targeted metabolomics across embryonic time reveals distinct onset of monoamine production **A–E.** Concentrations of five monoamines measured by targeted LC–MS/MS across seven developmental windows (0–3, 3–6, 6–9, 9–12, 12–15, 15–18, 18–22 h AEL). Each dot is one biological replicate (≥6 per time point); colour encodes the time window. The black line connects the mean of replicates at each time. The y-axis shows log_10_(normalised peak area + 1). The red dashed line marks the empirical detection threshold for each analyte. **A. Serotonin** and **B. Dopamine:** low/near-baseline early, followed by a sharp rise beginning ∼12–15 h, consistent with the transcriptional induction of biosynthetic enzymes. **C. Octopamine:** largely under the detection threshold. **D. Histamine** and **E. Tyramine**: detectable throughout development with gradual increases. Together, these profiles indicate that the functional availability of serotonin and dopamine lags their transcriptional commitment, whereas histamine and tyramine are present earlier and more continuously.

Together, these findings indicate that transcriptional programs broadly predict the functional potential of monoaminergic neurons, but the timing and extent of neurotransmitter production are modulated by subtype-specific biochemical properties and/or regulatory constraints.

### Conserved transcriptional dynamics underpin monoaminergic neuron development in *Drosophila* and zebrafish

Monoaminergic neurons are widely distributed across bilaterian animals, yet whether they arise from a conserved developmental program remains unclear. Our findings in *Drosophila* suggest that a specific set of TFs are activated during monoaminergic neuron differentiation, providing an opportunity to test the evolutionary conservation of this regulatory program. To address this question, we reanalysed published developmental single-cell RNA-seq data from *Danio rerio* and sea urchin, *Strongylocentrotus purpuratus*. For *Danio rerio*, we reprocessed a developmental time series spanning embryonic to larval stages (10 hours post-fertilisation [hpf] 232 to 10 days post-fertilisation [dpf]; Lange *et al*., 2024). For *S. purpuratus*, we merged two complementary datasets, covering the 8-cell embryo through the 72 hpf larval stages (Foster, Oulhen and Wessel, 2020; Perillo *et al*., 2020). Central nervous system cell types were annotated using orthologous marker genes applied previously in *Drosophila* (**Figure 2D**.) and identified monoaminergic neuron populations based on expression of canonical biosynthetic genes (**Figure S12A-B.**).

Applying the same differential-expression (DE) criteria used in fly (adjusted p < 0.05; |log₂FC| ≥ 0.25), we identified transcription factors (TFs) enriched in zebrafish and sea urchin monoaminergic clusters analysed independently (**Figure 6A**.; **Table S3.**). We analysed clusters separately, rather than collapsing them into a single monoaminergic group, to gather an inclusive list of TFs that define monoaminergic identity across subtypes. To enable cross-species comparisons despite lineage-specific duplications, genes were mapped to bilaterian-level eggNOG orthogroups. Orthologs of key fly regulators were recovered in vertebrates, including *lmx1a* paralogs (*lmx1a*, *lmx1al*, *lmx1ba*). In total, 15 TF orthogroups were differentially expressed in both fly and zebrafish monoaminergic neurons, and 16 TF orthogroups were shared between fly and sea urchin (**Figure 6C**.; **Table S4.**), defining a set of putative conserved regulators. Notably, six TF orthogroups—*CG8301*, *Lmx1a*/*CG4328*, *Hr3*, *Eip78C*, *net*, and *Pdp1*—were shared across all three species comparisons, underscoring a potentially conserved core program for monoaminergic identity. The partial overlap of the remaining TFs suggests that, while a conserved regulatory core might exist, additional factors have diverged or been co-opted in lineage-specific contexts

**Figure 6.**
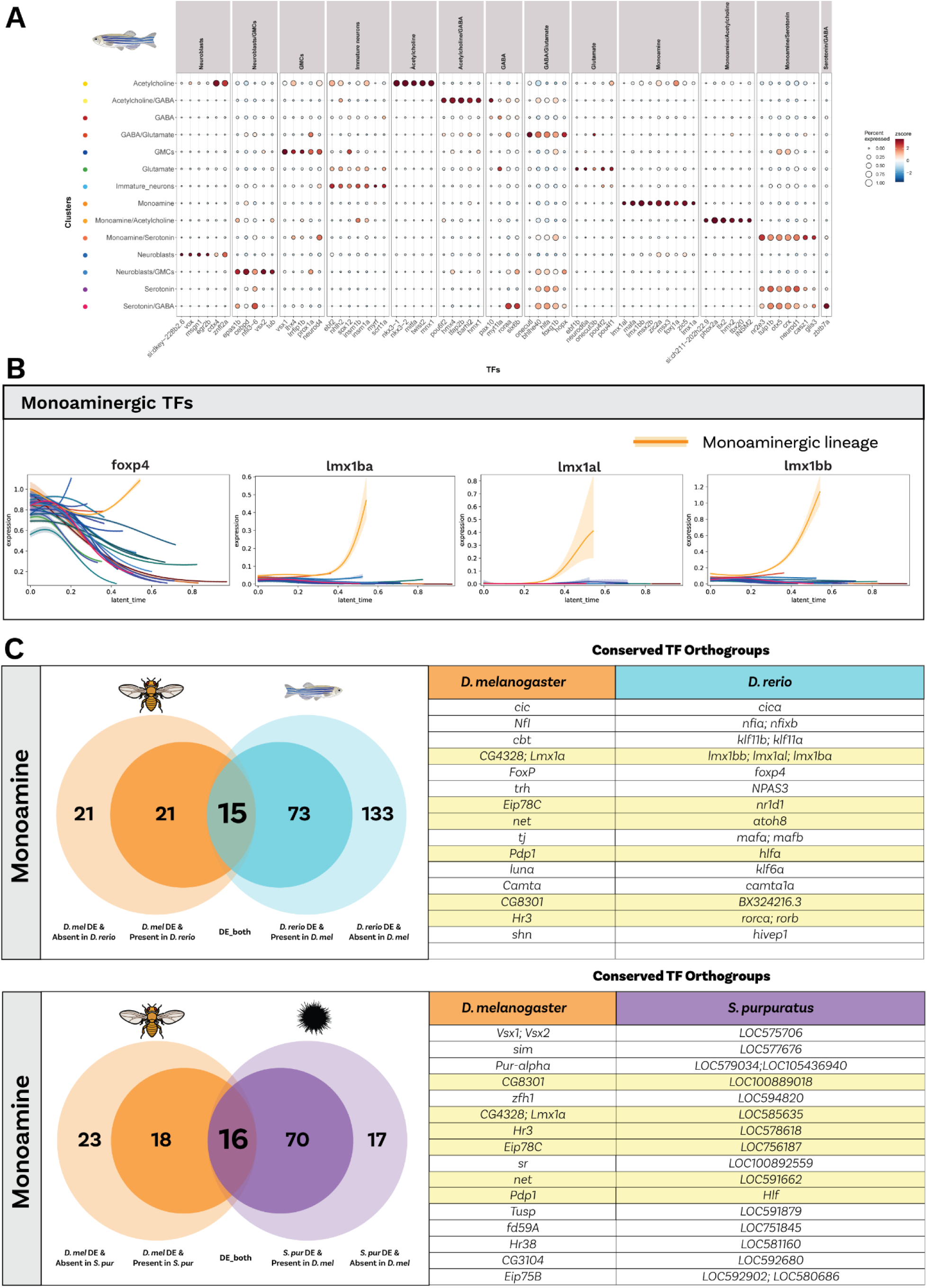
Cross-species conservation of monoaminergic TF programs **A. Differentially expressed TFs in *D. rerio* neurogenic subtypes**. Dot plot of differentially expressed TFs across zebrafish neurogenic subtypes. Columns are grouped by transmitter/state classes (Acetylcholine, Glutamate, GABA, Serotonin, Dopamine, Monoamine; plus, developmental states such as Neuroblasts, GMCs, Immature/Mature neurons). Dot size shows the fraction of cells in the subtype expressing the TF; colour shows the z-scored average expression. **B. Monoaminergic TF dynamics.** Smoothed latent-time expression trajectories for TFs associated with monoaminergic fate (examples shown: *foxp4*, *lmx1ba*, *lmx1a1*, *lmx1a*). Curves are plotted for the major neuronal lineages; the monoaminergic lineage is highlighted in orange with a light-orange confidence band, illustrating late-latent-time up-regulation of these TFs along the monoaminergic trajectory. **C. Orthogroup-level conservation analysis.** Venn diagram summarising Bilateria eggNOG orthogroups containing TFs that are differentially expressed in monoaminergic clusters in *D. melanogaster* and *D. rerio* and *D. melanogaster* and *S. purpuratus* (criteria as in Methods: adjusted p < 0.05; |log2FC| ≥ 0.25). Left/right circles denote species-specific DE orthogroups; the intersection denotes orthogroups DE in both species. The table lists representative conserved TF orthogroups with the corresponding gene names in each species. Together, these panels show that a core set of TF orthogroups is shared across insects and vertebrates while each lineage also exhibits species-specific components of the monoaminergic program.

To test whether this conservation was unique to monoaminergic neurons, we repeated the orthogroup-level analysis between *Drosophila* and zebrafish for other neurotransmitter classes (Acetylcholine, GABA, Glutamate; **Figure S13.**). The number of shared DE TF orthogroups was of similar magnitude across classes, despite vastly different neuronal population sizes, supporting a model in which each transmitter lineage may share a small, conserved set of regulators that is then diversified by lineage- and species-specific additions.

We next asked whether not only TF identity but also their temporal activation is conserved. Using the comparable time-resolved zebrafish dataset, we applied RNA velocity and CellRank to reconstruct differentiation dynamics. Several orthologous TFs, including *lmx1* paralogs, showed late upregulation along the zebrafish monoaminergic trajectory, closely mirroring the dynamics observed in *Drosophila* (**Figure 6B**.). Thus, some of the regulatory content and timing of activation are aligned between insects and vertebrates, at least in these representative species.

Together, these findings support a model in which monoaminergic neurons emerge from a potential conserved transcriptional program, with shared regulators, such as *Lmx1a*, but critically has undergone significant species-specific elaborations.

## Discussion

The extent to which homologous cell types are specified by conserved developmental and transcriptional programs remains a central question in evolutionary developmental biology. Monoaminergic neurons, which play fundamental roles in neurotransmission across bilaterians, offer a powerful model for addressing this issue. Previous comparative transcriptomic analyses have suggested that monoaminergic cell identity is defined by a shared set of transcription factors, indicating an origin in the bilaterian stem lineage (Goulty *et al*., 2025). Yet, whether the developmental processes that give rise to these neurons are likewise conserved—or instead represent cases of convergent evolution—has remained unresolved. Additionally, by determining whether the developmental program underlying these neurons is shared across species, we further clarify the hypothesis of a conserved evolutionary origin. In this study, we characterise the transcriptional program governing monoaminergic neuron development in *Drosophila* embryos and reveal that key regulatory components are shared with vertebrate and deuterostome systems. Our findings suggest that monoaminergic neurons are specified by an evolutionarily conserved developmental framework that emerged early in bilaterian evolution.

Within *Drosophila*, we identify a regulatory program involving *fd59A*, *dmrt99B*, *Vsx1*, *Vsx2*, and *Lmx1a*. These TFs are selectively expressed in the monoaminergic lineage and become active around stage 12, coincident with the onset of biosynthetic enzyme expression (*ple*, *Tdc2*, *Tbh*, *Vmat*). Spatial validation by HCR confirmed the sequential activation of these genes and showed that TF induction precedes biosynthetic gene expression, supporting their role as primary regulators of monoaminergic fate. This organisation mirrors vertebrate serotonergic development, where Lmx1a/b and Pet-1/Fev initiate comparable programs (Ding *et al*., 2003; Hendricks *et al*., 2003; Flames and Hobert, 2011), pointing to deep evolutionary parallels.

Although biosynthetic gene expression initiates around stage 12, metabolomic profiling reveals pronounced differences in monoamine accumulation. Dopamine and serotonin both increase after stage 12, yet dopamine accumulates more rapidly—likely reflecting the higher catalytic efficiency of Dopa decarboxylase (Ddc) for L-3,4- dihydroxyphenylalanine (L-DOPA) than for 5-hydroxytryptophan (5-HTP), the respective intermediates in the dopamine and serotonin biosynthetic pathways (Montioli and Borri Voltattorni, 2021). In contrast, octopamine remains undetectable despite enzyme expression, suggesting either technical limitations or additional biological constraints such as substrate availability, enzymatic kinetics, or post-transcriptional control. These findings indicate that transcriptional activation alone is insufficient for transmitter synthesis, pointing to additional regulatory layers that modulate monoamine output. However, these interpretations remain constrained by the detection sensitivity and chemical properties of the specific metabolites analysed. Our comparative analysis of monoaminergic neuron development in *Drosophila*, zebrafish, and sea urchins reveals a deeply conserved transcriptional program, underscoring a shared regulatory logic that likely emerged early in bilaterian evolution. The identification of six conserved TF orthogroups—*CG8301*, *Lmx1a*/*CG4328*, *Hr3*, *Eip78C*, *net*, and *Pdp1*—establishes a core genetic program for specifying monoaminergic identity. This finding supports the hypothesis that these neurons represent potentially homologous cell types that originated early in bilaterian evolution. Core regulatory modules specifying neuronal identity appear to have been established in the bilaterian ancestor and maintained across deep evolutionary time (Arendt, 2005; Arendt *et al*., 2019; Rapti, 2022; Sachkova, Modepalli and Kittelmann, 2025). For instance, Lmx1a has a well-documented and critical role in the development of dopaminergic neurons in vertebrates, where it regulates key effector genes such as tyrosine hydroxylase (*Th*) and the dopamine transporter (*DAT*) to ensure proper differentiation and maturation (Cai *et al*., 2009; Doucet-Beaupré, Ang and Lévesque, 2015). This role has not previously been demonstrated in non-vertebrates, suggesting that *Lmx1a* may represent an ancient regulator of monoaminergic fate specification—although convergent evolution of similar transcriptional strategies cannot be excluded.

Importantly, this conservation is not absolute. We also observed substantial regulatory divergence, with several TFs associated with monoaminergic differentiation being unique to specific lineages. This pattern may reflect a dual-layered evolutionary process—an ancient, conserved core program governing fate commitment, complemented by lineage-specific regulators that refine neuronal subtype identity (Hobert, 2008; Arendt *et al*., 2019). Alternatively, the conserved TF set itself may specify distinct monoaminergic identities across species, while divergent TFs act as complementary components that integrate these core programs into broader gene-regulatory networks, coordinating cross-talk and aligning specification with species-specific neural architectures (Flames and Hobert, 2011; Rapti, 2022). While some apparent differences may arise from technical factors (e.g., sequencing depth, detection sensitivity, or annotation), others likely represent genuine regulatory divergence. Given the substantial variation in neuroanatomical organisation among insects, vertebrates, and echinoderms, such divergence may reflect adaptive remodelling of the regulatory landscape to distinct neural architectures.

The temporal dynamics of conserved TFs suggest that these genes govern monoaminergic fate commitment, whereas lineage-specific TFs modulate neuronal subtype identity in a species-dependent manner. The hierarchical relationships between conserved and lineage-specific TFs remain to be elucidated. Determining whether these factors operate through feedforward or feedback regulatory loops—a recurrent feature of GRN architecture that stabilises neuronal identity—will be crucial (Flames and Hobert, 2011; Peter and Davidson, 2011). Understanding how conserved GRNs are modified to generate lineage-specific phenotypes will be essential for uncovering the general principles that govern cell-type diversification across evolution. Our mass-spectrometry data indicate a partial decoupling between monoamine abundance and biosynthetic enzyme expression. While enzyme kinetics may account for some of the observed differences in transmitter accumulation, it remains unclear whether such decoupling is a general feature of monoaminergic neurons across taxa. Moreover, the functions of early monoamine signalling—potentially enabled by precocious receptor expression (e.g., 5-HT2A)—during embryogenesis remain unresolved.

Integrating transcriptomic, metabolomic, and spatial data across additional bilaterian taxa will be important for testing the universality of these patterns. Comparative perturbation experiments—targeting conserved TFs such as Lmx1a or Hr3 in non-model systems—could determine whether shared transcriptional logic indeed translates into conserved functional outcomes.

In conclusion, our study uncovers a conserved temporal program for monoaminergic neuron specification, in which early transcription factor activation orchestrates biosynthetic gene expression to align fate determination with functional maturation. By linking transcriptional programs, transmitter synthesis, and receptor deployment across species, our work provides a framework for understanding how conserved and divergent GRNs generate neuromodulatory diversity and how neuronal fate and function are developmentally coupled.

## Methods

### Fly husbandry

All *Drosophila melanogaster* stocks were maintained on standard cornmeal–yeast– agar medium at 18°C or 25°C, with relative humidity held at approximately 40%. We employed a transgenic SoxN–sfGFP line (VDRC #318062), which was reared continuously at 25°C (Sarov *et al*., 2016).

### Embryo collection

Adult flies were transferred into population cages fitted with agar plates prepared from mango juices, each supplemented with fresh yeast paste to boost fecundity. To eliminate circadian-driven fluctuations in gene expression, cages were kept in constant darkness (DD) beginning immediately upon transfer (Wijnen *et al*., 2006). Following a 48-hour acclimation under DD, embryo collections was commenced, ensuring that all developmental stages progressed without light-induced transcriptional variability.

Embryos were collected over eight successive, overlapping time windows covering the full span of embryonic development; between 0-22 hours after egg laying (AEL). Collections were performed in biological duplicate to increase data reliability and resolution of transitional stages. Agar plates were replaced regularly to maintain optimal oviposition conditions, and collected embryos remained in the same DD incubator for precise aging.

Given the necessity of preserving RNA integrity for 10X Genomics single-cell RNA-sequencing, we avoided formaldehyde-based fixatives. Instead, we applied dithio-bis(succinimidyl propionate) (DSP; Sigma-Aldrich), a reversible cross-linker compatible with downstream library preparation. A 50 mg/mL DSP stock was made in DMSO, then diluted by adding 5 mL Dulbecco’s PBS (DPBS; Gibco) to the 100 µL stock, followed by sterilisation through a 0.2 µm syringe filter. Collected embryos were fixed in this working solution, preserving stage-specific transcriptomes and enabling reliable reversal of cross-links prior to sequencing.

### Embryo Harvesting, Dechorionation, and Fixation

Collected embryos were gently dislodged from agar plates by rinsing with distilled water and transferred into a 30 µm nylon mesh sieve using a fine paintbrush. Any residual yeast paste was washed away under a gentle stream of distilled water to minimise contamination. Dechorionation was performed by submerging the embryos in 50% bleach (Arco Essentials) for 3 minutes, then immediately rinsing them thoroughly with distilled water until all bleach was removed.

For fixation, equal volumes of the DSP working solution and heptane (Fisher Scientific) were combined in a glass scintillation vial. Dechorionated embryos were added to this biphasic mixture and incubated on a gyroscopic rocker at room temperature for 1 hour, ensuring gentle but continuous agitation. After fixation, the lower aqueous phase was discarded, and cold methanol (VWR) was added to the heptane layer to initiate devitellinisation. Embryos were vortexed for 2 minutes to facilitate complete removal of the vitelline membrane, then allowed to settle to the bottom of the vial. Finally, embryos were transferred to 1.5 mL Eppendorf tubes and washed three times in fresh methanol. Fixed embryos were stored at –20°C until further processing.

### Immunofluorescence

All immunostaining was performed at 4°C under stringent RNase-free conditions: Protector RNase Inhibitor (Roche, RNAINH-RO) was added to all buffers at 5 U/mL during brief washes and 20 U/mL for all longer incubations, and both bovine serum albumin (BSA; Gibco) and normal goat serum (NGS; Vector Laboratories) were heat-inactivated at 60°C for 30 minutes to eliminate residual RNases. Prior to antibody staining, DSP–methanol–fixed embryos were gradually rehydrated through an ethanol series into PBT (PBS + 0.1% Triton X-100; Sigma-Aldrich T8787) and then washed three times for 10 minutes each in PBT containing RNase inhibitor, with gentle rocking to ensure complete removal of the methanol fixative. To block non-specific binding, embryos were incubated overnight at 4°C in PBT supplemented with 1% heat-inactivated BSA and 2% heat-inactivated NGS. The embryos were then transferred into fresh blocking buffer (PBT + inhibitor, 1% BSA, 2% NGS) containing primary antibodies: rabbit α-GFP (1:100; Invitrogen) to label SoxN–sfGFP⁺ neurogenic cells, and rat α-Elav (1:100; DSHB) to detect post-mitotic neurons. After overnight incubation at 4°C with gentle agitation, excess primary antibody was removed by three 10-minute washes in PBT. Secondary detection was performed by incubating embryos overnight in the same blocking buffer supplemented with goat α-rabbit Alexa Fluor 488 (1:500; Invitrogen), mouse α-rat PE (1:500; Invitrogen), and the nuclear counterstain DRAQ5 (4 µL/mL; Thermo Scientific). Finally, stained embryos were washed three times in PBT (10 minutes each) and stored at 4 °C in PBT until mounting and imaging.

### Microscopy validation and processing

Images were acquired using a Zeiss LSM 980 Airyscan 2 microscope in Airyscan super-resolution mode, with the default settings applied for Airyscan image processing. A Plan-Apochromat 10x/0.45 or Plan-Apochromat 20x/0.8 objective was employed for image acquisition (University of Leicester Advanced Imaging Facility). Post-processing was performed using a custom FIJI (Schindelin *et al*., 2012) macro for orienting the embryos along anatomical axes (<u>Embryo Processing IF Script</u>).

### Single-cell dissociation

Embryos were transferred into an ice-cold 15 mL Dounce homogeniser in PBS containing Protector RNase Inhibitor, and all subsequent steps were performed at 4°C to preserve RNA integrity. After three gentle washes with cold Dulbecco’s PBS (DPBS), the supernatant was removed and replaced with PBS supplemented with 0.04% BSA (PBS/BSA). Embryos were then mechanically disrupted by 10–15 strokes with the tight-fitting pestle, and the homogenate was collected into a fresh tube. To maximise cell recovery, the homogeniser was rinsed with additional PBS/BSA, and the combined suspension was centrifuged at 3,000 × g for 3 minutes. The supernatant was discarded, and the pellet was resuspended in PBS/BSA.

To remove large debris, the resuspended cells were centrifuged at 40 × g for 3 minutes, after which the supernatant—containing single cells—was transferred to a new tube and spun at 1,000 × g for 3 minutes. The resulting pellet was gently triturated in PBS/BSA by passing the suspension through a 22-gauge needle with 20 strokes to dissociate any remaining aggregates, then filtered through a 100 µm nylon mesh. A final spin at 1,000 × g for 3 minutes pelleted the cleaned single-cell suspension, which was resuspended in PBS/BSA and kept on ice prior to fluorescence-activated cell sorting (FACS). Parallel control samples—unstained; stained with only primary or only secondary antibodies; and combinations thereof with DRAQ5 nuclear stain—were prepared and processed identically for gating and compensation during FACS analysis.

### FACS

Cells were sorted on a Beckman Coulter MoFlo Astrios cell sorter (Nottingham) equipped with a 100 µm nozzle, operating in “Purify” precision mode with a 1–2 drop envelope to maximise purity of collected fractions. Single-cell suspensions were maintained at 4°C throughout sorting by housing the entire flow chamber in a water-cooled environment, thereby minimising heat-induced stress and preserving cell integrity. Samples were deposited into a two-way tube holder pre-chilled on ice.

Gating strategies were defined using unstained and single-stained controls to eliminate autofluorescent events. We applied broad gates to capture both strongly and weakly fluorescent populations, acknowledging the continuum of marker expression. SoxN⁺ and Elav⁺ cells—representing early neurogenic and post-mitotic neuronal cohorts, respectively—were collected together. This inclusive approach ensured recovery of the full spectrum of developing neural cells.

### Reverse cross-linking

After FACS, the cells were subjected to a reverse cross-linking step to remove the DSP fixative and prepare them for subsequent library preparation. The sorted cells were first centrifuged at 4°C at 1,000 x g for 3 minutes, and the supernatant was discarded. The cell pellet was then resuspended in PBS/BSA solution and incubated with 50 mM dithiothreitol (DTT, Sigma-Aldrich) at 37°C for 30 minutes to break the cross-links formed by DSP. Following incubation, the cells were centrifuged twice at 4°C at 1,000 x g for 3 minutes to thoroughly remove any residual DTT.

## 10X Genomics Library Preparation

The library preparation was conducted using the 10X Genomics Chromium Next GEM Single Cell 3’ Reagent Kit v3.1 (10X Genomics). Prior to the preparation, cell concentration was assessed with a Luna FX7 Cell Counter, and cells were adjusted to a concentration of approximately 500-1500 cells/µl. Approximately 8000 cells per sample were used to generate single-cell gel beads in emulsion (GEMs), targeting approximately 5000 single-cell GEMs per sample. GEM emulsions were created with the 10X Chromium Controller, followed by complementary DNA (cDNA) synthesis and library preparation according to the manufacturer’s protocol. cDNA amplification was performed for 12 cycles, and library amplification during the PCR indexing stage was similarly carried out for 12 cycles using the Dual Index Kit TT Set A (10X Genomics).

Quantification of cDNA was performed using Qubit dsDNA HS assays (Invitrogen) and D5000 HS ScreenTape (Agilent) on an Agilent Technologies TapeStation 4200, while library quantification was conducted with Qubit dsDNA HS assays and D1000 ScreenTape on the same platform. Libraries were prepared following the manufacturer’s guidelines and were subsequently sequenced externally by Novogene on the Illumina NovaSeq X Plus platform, utilising 150 bp paired-end reads.

### HCR Validation

Custom probe sets targeting mRNAs of key monoaminergic pathway enzymes, receptors, and TFs were designed from the longest isoform FlyBase annotations and synthesised by Molecular Instruments. Briefly, embryos were first equilibrated in hybridisation buffer (v3.1, Molecular Instruments) at 37°C for 30 min to block non-specific sites, then incubated overnight (12–16 h) at 37°C with 4 pmol of each probe set in fresh hybridisation buffer, following the manufacturer guidelines. Following this, unbound probes washed, and fluorophore-conjugated HCR hairpins (Molecular Instruments) were snap-cooled and applied to embryos in amplification buffer, then incubated for 12–16 h at room temperature in the dark. Post-amplification, embryos were washed to remove excess hairpins.

Finally, embryos were counterstained with DAPI (1:1000, Sigma-Aldrich) during the last wash, mounted in antifade medium (Vectashield; Vector Laboratories), and imaged by confocal microscopy and processed using a custom macro script in FIJI (Embryo Processing HCR Script).

### Targeted Metabolomics

Embryos from seven consecutive developmental windows were rinsed thoroughly in distilled water to remove yeast, excess liquid discarded, weighed, and flash-frozen in liquid nitrogen. Samples were shipped on dry ice to the EMBL Metabolomics Core Facility (Heidelberg, Germany), where Bernhard Drotleff performed all downstream processing.

Each frozen sample (75 mg/mL) was extracted on dry ice by adding 80% acetonitrile (with internal standards and 1% formic acid), then homogenised using 1 mm zirconia beads in a bead-beater (FastPrep-24) at 6.0 m/s (5 × 30 s, 5 min rest). After incubation at –20 °C for 20 min, lysates were vortexed and centrifuged (15 000 × g, 10 min, 4 °C). Supernatants were transferred to glass vials and analysed within 2 h.

LC–MS/MS was carried out on an Agilent Infinity II BioLC coupled to a Sciex QTRAP 6500+ (positive ESI). Chromatography used an Atlantis Premier BEH Z-HILIC column (2.1 × 100 mm, 1.7 µm) at 40 °C, with mobile phases A (90:10 H₂O:ACN) and B (98:2 ACN:H₂O), both buffered with 25 mM ammonium formate (pH 3.0). The 13 min gradient (0–1 min 100% B; 1–8 min to 60% B; 8–10 min 60% B; 10–10.5 min to 100% B; 10.5–13 min 100% B) ran at 0.3 mL/min, with 5 µL injections at 4 °C. Ion source parameters were: curtain gas 40 psi, nebulizer gas 40 psi, heater gas 40 psi, ion spray voltage 5 kV, source temperature 400 °C, medium collision gas. Data were acquired in scheduled MRM mode (90 s windows, 600 ms cycle). Samples were analysed in randomised order, with pooled QCs for system equilibration and every fifth sample to monitor performance; a processed blank defined background. Raw peak areas were extracted and processed in SciexOS. Data analysis and visualisation was done in R using custom scripts (<u>Metabolomics R Script</u>).

### scRNA-seq Analysis: Data alignment

Raw sequencing data were processed using Cell Ranger count (v9.0.1, 10X Genomics). Reads were aligned to a custom *Drosophila melanogaster* reference (Ensembl release BDGP6.46.110) into which the GFP transgene sequence had been incorporated. Cell Ranger generated UMI-collapsed gene–cell count matrices for each library.

### scRNA-seq Analysis: Preprocessing and Quality Control

Downstream quality control and preprocessing were carried out in Seurat (v5.2.0; full pipeline available at <u>GitHub</u> page). Briefly, cells with fewer than 800 or more than 100,000 UMIs, fewer than 200 or more than 8,000 detected genes, or mitochondrial transcript content exceeding 8–10% were excluded. After log-normalisation and identification of highly variable features, JackStraw analysis was used to select significant principal components for dimensionality reduction. Uniform Manifold Approximation and Projection (UMAP) was then performed on these PCs. To remove artifactual multiplets, DoubletFinder (v2.0.6, McGinnis, Murrow and Gartner, 2019) was applied with an expected doublet rate of 3.9%, consistent with 10X Genomics loading recommendations.

### scRNA-seq Analysis: Integration and Cluster Annotation

Biological replicates from each developmental time point were integrated using Harmony (v1.2.3, Korsunsky *et al*., 2019) to correct batch effects while preserving biological variation. Clusters were initially defined in the integrated UMAP space and then annotated following a modified version of the Calderon *et al*. (2022) pipeline: marker genes for each cluster, selected as the top differentially expressed features with adjusted P < 0.05, were compared against curated spatial expression terms from the Berkeley Drosophila Genome Project (BDGP) using Fisher’s exact test to identify over-represented *in situ* annotations (Tomancak *et al*., 2002, 2007; Hammonds *et al*., 2013; Calderon *et al*., 2022). After Bonferroni correction, significant enrichment terms were manually reviewed and consolidated to assign each cluster to a defined neurogenic or neuronal identity. This combined data-driven and expert-curated approach ensured robust annotation of cell states across *Drosophila* embryonic neurogenesis.

To preserve the continuum of developmental variation, cells from all time points were merged without additional batch-correction; standard integration algorithms tended to over-correct and attenuate genuine developmental signals. The merged dataset was then reclustered in Seurat, and each resulting cluster was assigned an identity based on the modal annotation of its constituent cells from the individual time- point analyses. This majority-vote approach ensured that clusters reflected the predominant cell state across development.

### scRNA-seq Analysis: Neuronal-Specific Annotation

We next focused on delineating specific neuronal lineages. Clusters corresponding to the progression from neuroblasts to mature neurons were reannotated using expression of subtype markers defined in Dillon *et al*. (2022) Peng *et al*. (2024). This approach yielded fine-grained, cell-type annotations that aligned with, and extended, the broader developmental ontology derived from BDGP enrichment annotation pipeline.

### scRNA-seq Analysis: Trajectory Inference via RNA Velocity

To infer cellular dynamics and developmental trajectories, Cell Ranger BAM files were reprocessed with Velocyto (v0.17.17, La Manno *et al*., 2018) to quantify spliced and unspliced transcripts, producing Loom matrices. Next, the Seurat-annotated object was converted to an AnnData structure using sccustomize (v3.0.1, Samuel Marsh *et al*., 2024), and merged with the velocity matrices preserving metadata, embeddings, and clustering labels. Downstream modeling was carried out in scVelo (v0.3.3, La Manno *et al*., 2018; Bergen *et al*., 2020) using the dynamical model: data were first filtered and normalised, moments were computed on the neighbourhood graph, and velocity vectors were estimated for each cell. Finally, velocity-stream embeddings were projected onto the UMAP embedding to visualise inferred developmental progressions.

### scRNA-seq Analysis: Subclustering and Lineage Dynamic Analysis

To resolve finer neuronal subtypes, we isolated cells from the maturing neuronal lineage and performed a second round of clustering in Seurat. Clusters were reannotated based on the expression of established subtype marker genes, and dot plots were generated to verify marker specificity. We then recalculated RNA-velocity on this subset in scVelo (v0.3.3, La Manno *et al*., 2018; Bergen *et al*., 2020), re-deriving the first- and second-order moments, velocity vectors, velocity-stream embedding, and latent-time projections. These velocity maps were overlaid on the UMAP to visualise intra-lineage transitions. Finally, CellRank was applied to the subclustered dataset—integrating connectivity, velocity, and CytoTRACE kernels—to infer both initial (progenitor) and terminal (mature neuron) fates and compute fate probabilities within the neuronal lineage (Reuter *et al*., 2018; Lange *et al*., 2022; Reuter, Klein and Lange, 2022; Weiler *et al*., 2024).

### scRNA-seq Analysis: Regulatory dynamics

To uncover TFs that drive subtype specification, we performed differential expression analysis restricted to TFs annotated in the AnimalTFDB for *Drosophila melanogaster* (Shen *et al*., 2023). Candidate TFs with significant up-regulation in each neuronal subtype were identified and ranked. Furthermore, leveraging the latent-time ordering from scVelo, we modelled the dynamic expression of all genes along developmental trajectories. We then refined this analysis using CellRank’s multi-kernel framework— combining connectivity, velocity, and CytoTRACE information—to pinpoint TFs whose expression kinetics most strongly correlate with the latent time. This layered approach highlighted key regulators of neuronal differentiation for subsequent validation using HCR.

### Comparative scRNAseq analysis: Dataset Acquisition and Preprocessing

To explore the evolutionary conservation of these regulators, we reanalysed published single-cell RNA-seq datasets from *Danio rerio* and *Strongylocentrotus purpuratus*. The zebrafish atlas spans 10 developmental stages from 10 hours post-fertilisation (hpf) to 10 days post-fertilisation (dpf) (Lange *et al*., 2024). The sea urchin dataset was generated from combining two datasets, comprising 10 time points from the 8-cell stage to the 72 hpf larval stage (Foster, Oulhen and Wessel, 2020; Perillo *et al*., 2020). For zebrafish, raw 10x Genomics sequencing data were reprocessed with Cell Ranger (v9.0.1; 10x Genomics) against the Ensembl *D. rerio* reference (Danio_rerio.GRCz11.114), retaining both spliced and unspliced reads. Resulting BAM files were converted to spliced/unspliced count matrices using Velocyto (v0.17.17, (La Manno *et al*., 2018), yielding loom files for downstream analysis. For sea urchin, we used the author-provided count matrices and accompanying metadata without remapping.

### Comparative scRNAseq analysis: Data Integration and Annotation

Rather than reclustering the full datasets, we focused on neural lineages. For zebrafish, central nervous system (CNS) populations were subset using the original annotations provided in the source atlas (https://zebrahub.sf.czbiohub.org/data). For sea urchin, we subset to neurogenic clusters based on the authors’ labels.

Cross-species re-annotation was performed with orthologous marker genes derived using eggNOG-based orthology mapping (described below), with the *Drosophila melanogaster* subclustering marker set as the anchor. For sea urchin, orthologues were called from the *S. purpuratus* proteome corresponding to the reference genome used for mapping (GCF_000002235.4; Spur_4.2). Zebrafish markers were referenced to Ensembl *D. rerio* GRCz11 (release 114). Monoaminergic neuron populations were identified by expression of the orthologous canonical biosynthetic enzymes and transporters.

### Comparative scRNAseq analysis: Trajectory and Dynamic Expression Analyses

RNA velocity was estimated using scVelo (Bergen *et al*., 2020) and lineage inference was refined with CellRank (v2) (Weiler *et al*., 2024) by integrating connectivity and velocity information. Latent-time ordering was used to model dynamic gene expression trajectories and identify TFs with temporal activation patterns correlating with neuronal differentiation.

### Comparative scRNAseq analysis: Differential Expression and Orthology Mapping

Candidate TFs enriched in each neuronal subtype were identified using the same thresholds applied to the *Drosophila* dataset (adjusted p < 0.05; |log2FC| ≥ 0.25). For cross-species comparison, we re-annotated the proteomes of zebrafish and *Drosophila* with eggNOG (v5.0, Huerta-Cepas *et al*., 2019), assigning genes to its hierarchical orthogroup framework. We then conducted all comparative analyses at the Bilateria-level orthogroups, which consolidate one-to-many and many-to-one relationships, enabling robust mapping of putative orthologs and fair TF comparisons across species.

### Comparative scRNAseq analysis: Comparative and Class-specific Analyses

Shared and species-specific regulators were identified by comparing differentially expressed TF orthogroups across *Drosophila* and zebrafish monoaminergic clusters. To contextualise these findings, the orthogroup-level analysis was extended to other neurotransmitter lineages (acetylcholine, GABA, glutamate).

## Data availability

All data supporting this study are available on Zenodo. Processed FACS data are deposited at https://doi.org/10.5281/zenodo.17449471, targeted metabolomics data at https://doi.org/10.5281/zenodo.17449991, and processed single-cell RNA-seq data at https://doi.org/10.5281/zenodo.17441737. All datasets are released under the MIT License. Analysis pipelines and scripts are available at https://github.com/cliftonlewis/2025_Drosophila_scRNAseq_EmbryoNeurogenesis_Monoamine.

## Author Contributions

R.F. conceived the project. C.L. performed the majority of sample preparation for scRNA-seq, HCR, and targeted metabolomics, and carried out the bioinformatics analyses. M.G. provided guidance on the bioinformatics analyses. A.W. assisted with optimization of HCR protocols. N.C. and D.O. contributed to FACS sorting. S.R. supported the initial project setup. R.P.Z. provided the initial protocols for targeted scRNA-seq in *Drosophila* embryos. J.S. contributed to project development. E.R. and C.P.K. supervised and supported the project. R.F. was the primary supervisor. C.L. and R.F. wrote and edited the manuscript.

## Acknowledgements

We gratefully acknowledge the support of the Royal Society through a University Research Fellowship (UF160226 and URF/R/221011) and a Research Grant (RGF\R1\181012) awarded to R.F. C.L. is supported by a BBSRC-MIBTP PhD Studentship (Grant number MIBTP2020: BB/T00746X/1). The authors wish to thank the technical and scientific staff of several core facilities for their invaluable support. We are especially grateful for the assistance received from the University of Leicester Advanced Imaging Facility (RRID: SCR_020967; BBSRC grant: BB/S019510/1) and specifically thank Dr. Kees Straatman for his expert microscopy advice. This research used the ALICE High Performance Computing Facility at the University of Leicester. We thank the Flow Cytometry Facility, particularly Dr. David Onion and Nicola Croxall, for their support with the Beckman Coulter MoFlo Astrios Flow Cytometer. We also thank the DEEP-SEQ Genomics Facility, specifically Dr. Christopher Moore and Dr. Nadine Holmes, for their expert assistance with single-cell genomics, including 10X Genomics support. Additionally, we are deeply appreciative of Dr. Bernhard Drotleff from the EMBL Metabolomics Facility for running the targeted metabolomics experiments. We would also like to thank Dr. Diego Calderon (University of California San Francisco) for help with adapting the cell type annotation pipeline. Finally, we would like to thank the Feuda Lab for their thoughtful proofreading and constructive feedback on the manuscript.

