## Supplementary Figures for "Evolutionarily conserved transcriptional regulators control monoaminergic neuron development"

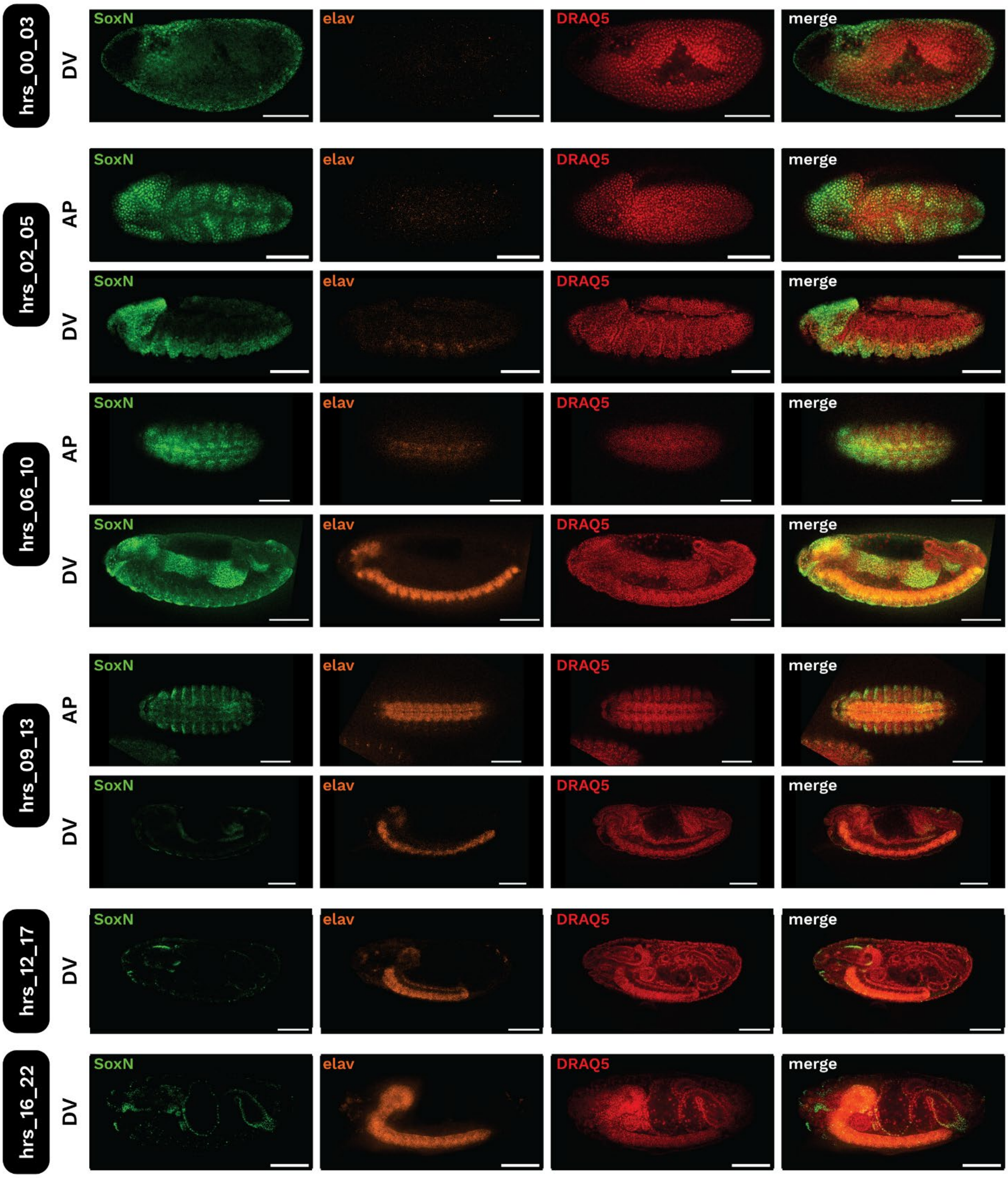


**Figure S1. Temporal expression dynamics of SoxN and Elav in *Drosophila* across embryogenesis.**

Confocal images showing immunofluorescence staining of *Drosophila* embryos at representative developmental windows using antibodies against SoxN (green, neural progenitor marker), Elav (orange, post-mitotic neuronal marker), and nuclear counterstain DRAQ5 (red). Embryos were imaged at six developmental stages: 0–3 hpf, 2–5 hpf, 6–10 hpf, 9–13 hpf, 12–17 hpf, and 16–22 hpf. Both anterior-posterior (AP) and dorsal-ventral (DV) views are shown where applicable. Early-stage embryos (0–5 hpf) display broad SoxN expression and minimal Elav signal. As development progresses, SoxN expression reduces, while Elav expression is progressively upregulated in differentiating neurons beginning ~6–10 hpf. By stage 13 (~12 hpf), Elav+ neurons are abundantly distributed throughout the VNC and brain lobes, while SoxN expression is minimal. Scale bars = 100 μm.

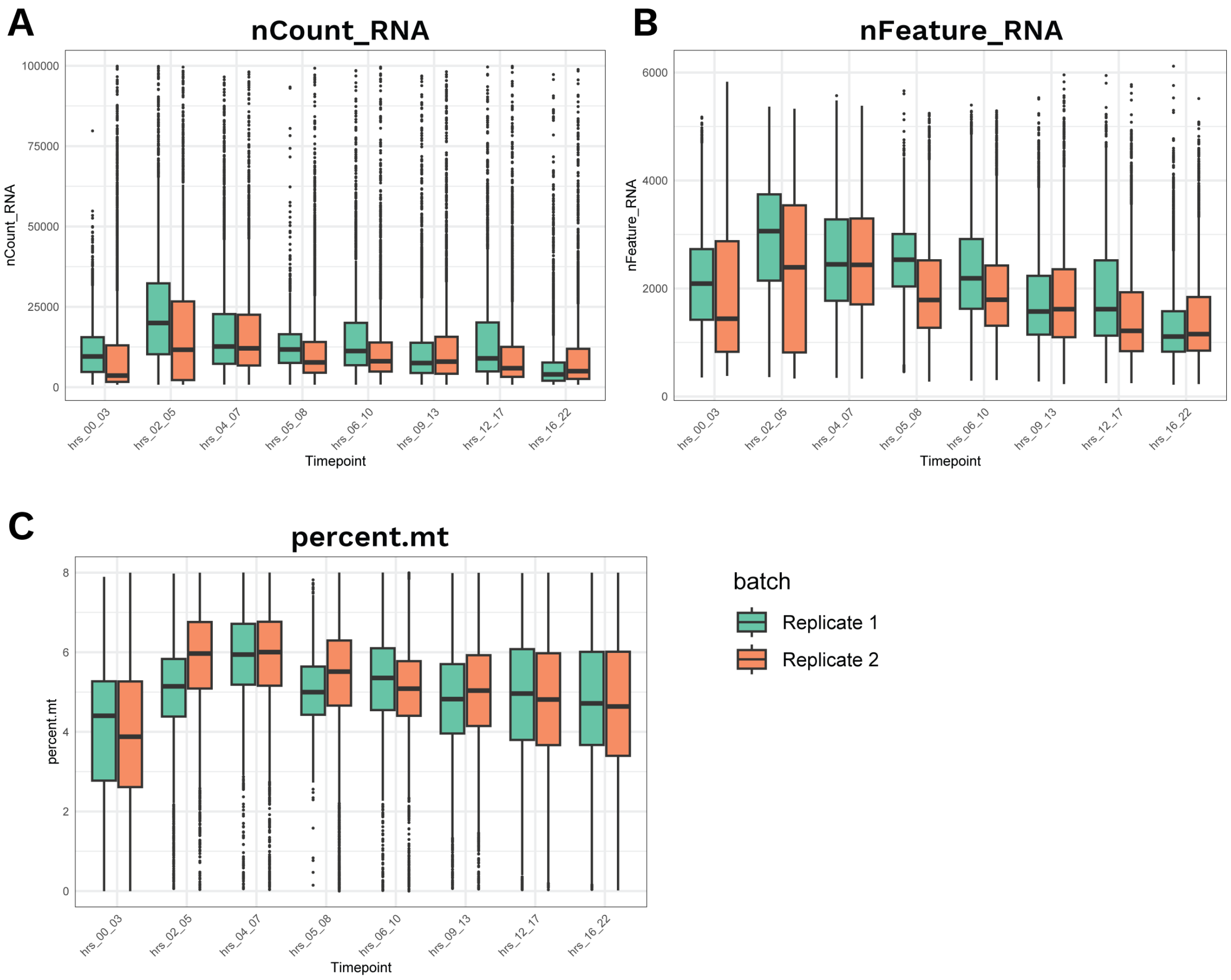


**Figure S2. Quality metrics by timepoint and replicate**

**A. UMI counts per cell** (*nCount_RNA*), **B. detected genes per cell** (*nFeature_RNA*), and **C. mitochondrial RNA fraction** (*percent.mt*). Boxplots show the median and IQR; whiskers extend 1.5×IQR; points are individual cells. Colors denote biological replicates (Replicate 1, Replicate 2).

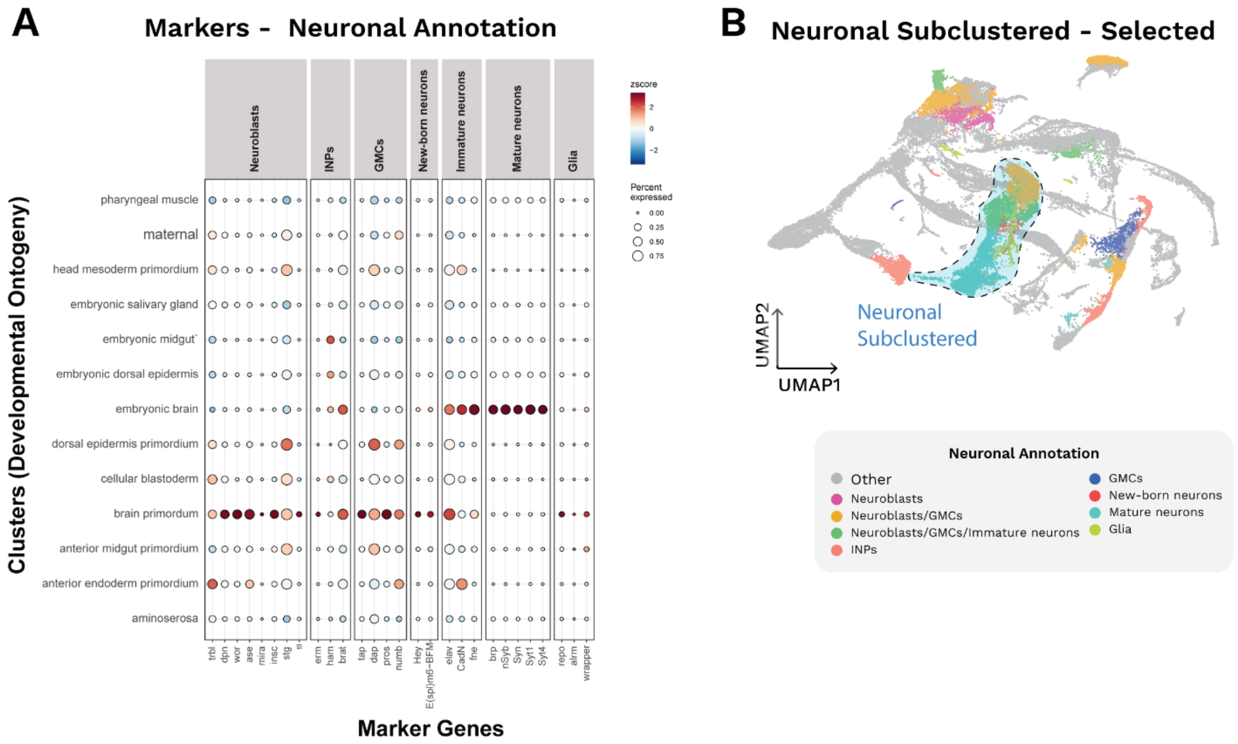


**Figure S3. Congruence between neuronal-state annotations and developmental ontogeny, and selection of the maturing neuronal lineage for subclustering.**

**A.** Dot plot comparing literature-curated neuronal markers with clusters annotated by developmental ontogeny. Rows list ontogeny-defined clusters (e.g., maternal, cellular blastoderm, brain primordium, embryonic brain, and non-neural tissues). Columns group genes by neuronal state: neuroblasts, intermediate neural progenitors (INPs), ganglion mother cells (GMCs), new-born neurons, immature neurons, mature neurons, and glia. Dot size indicates the fraction of cells in a cluster expressing a given gene; colour reflects z-scored average expression. **B.** UMAP of the neuronal atlas with the **'Neuronal subclustered'** selection outlined in blue. These selected clusters correspond to the maturing neuronal lineage and were carried forward for downstream subclustering.

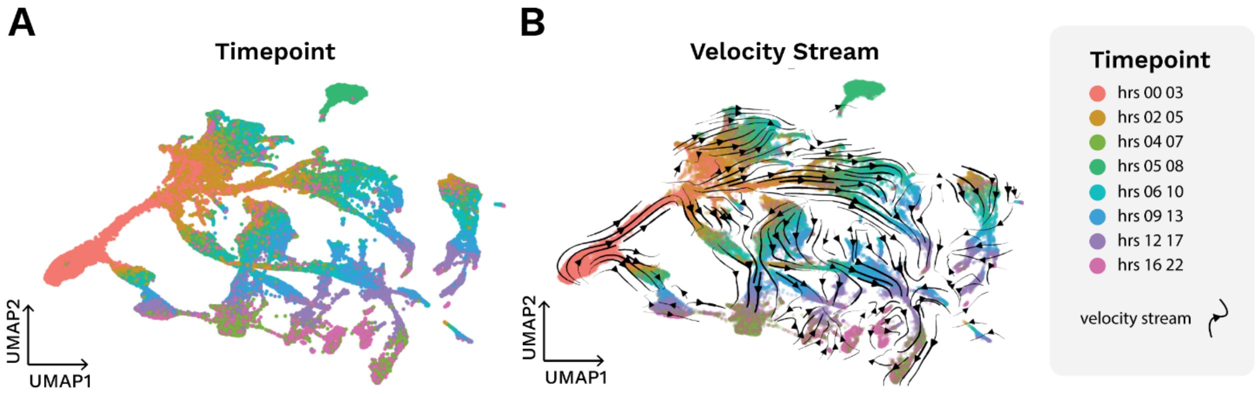


**Figure S4. Concordance of timepoint annotations with RNA velocity across the full dataset**

**A.** UMAP plot coloured by timepoint. **B.** Same UMAP embedding with velocity streamlines overlaid. Streamlines run along the same manifold and point from early to late timepoints, indicating that sampling times and inferred dynamical flow agree.

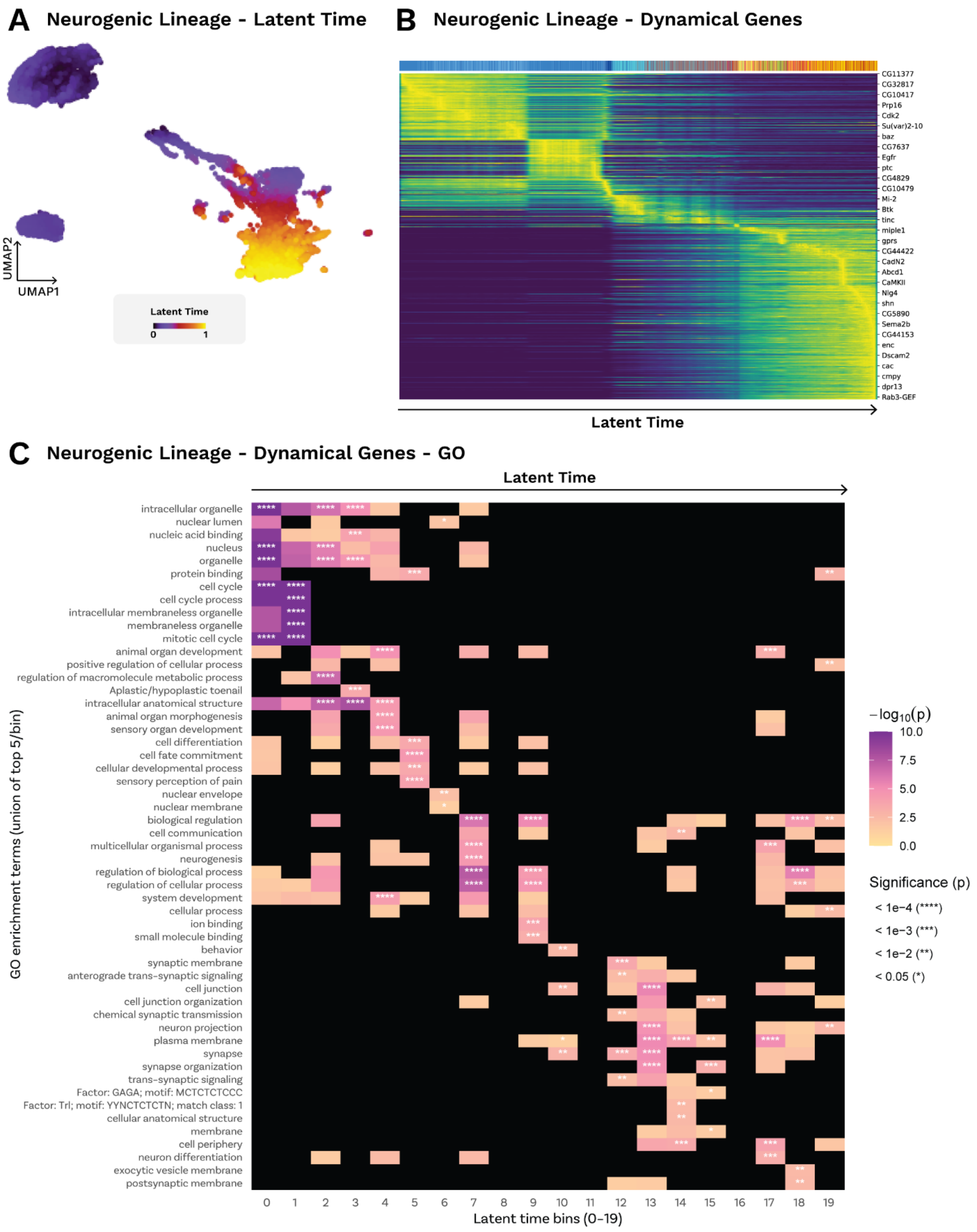


**Figure S5. Neurogenic lineage – Latent time, dynamical genes, and gene ontology across the trajectory**

**A. Latent time**. UMAP of the neurogenic lineage subset coloured by scVelo latent time (0→1). The arrow denotes the inferred developmental progression. **B. Dynamical genes.** Heatmap of top scVelo dynamical genes, with cells ordered by latent time and genes ordered by their peak latent-time of expression. Values are z-scored per gene;

brighter colours indicate higher expression. **C. Gene Ontology across latent time.** Dynamical genes were assigned to one of 20 equal latent-time bins by their peak time and tested for GO enrichment (g:Profiler; *D. melanogaster*; GO:BP and GO:MF). Tiles show enrichment magnitude (fill = $-\log10$ p); white dots mark FDR-significant terms (BH-adjusted $p \leq 0.05$). Terms are displayed at the bin where enrichment peaks, revealing a trajectory from early organelle/biogenesis and cell-cycle programs to late neuronal differentiation, synaptic specification, and axon/neurite functions.

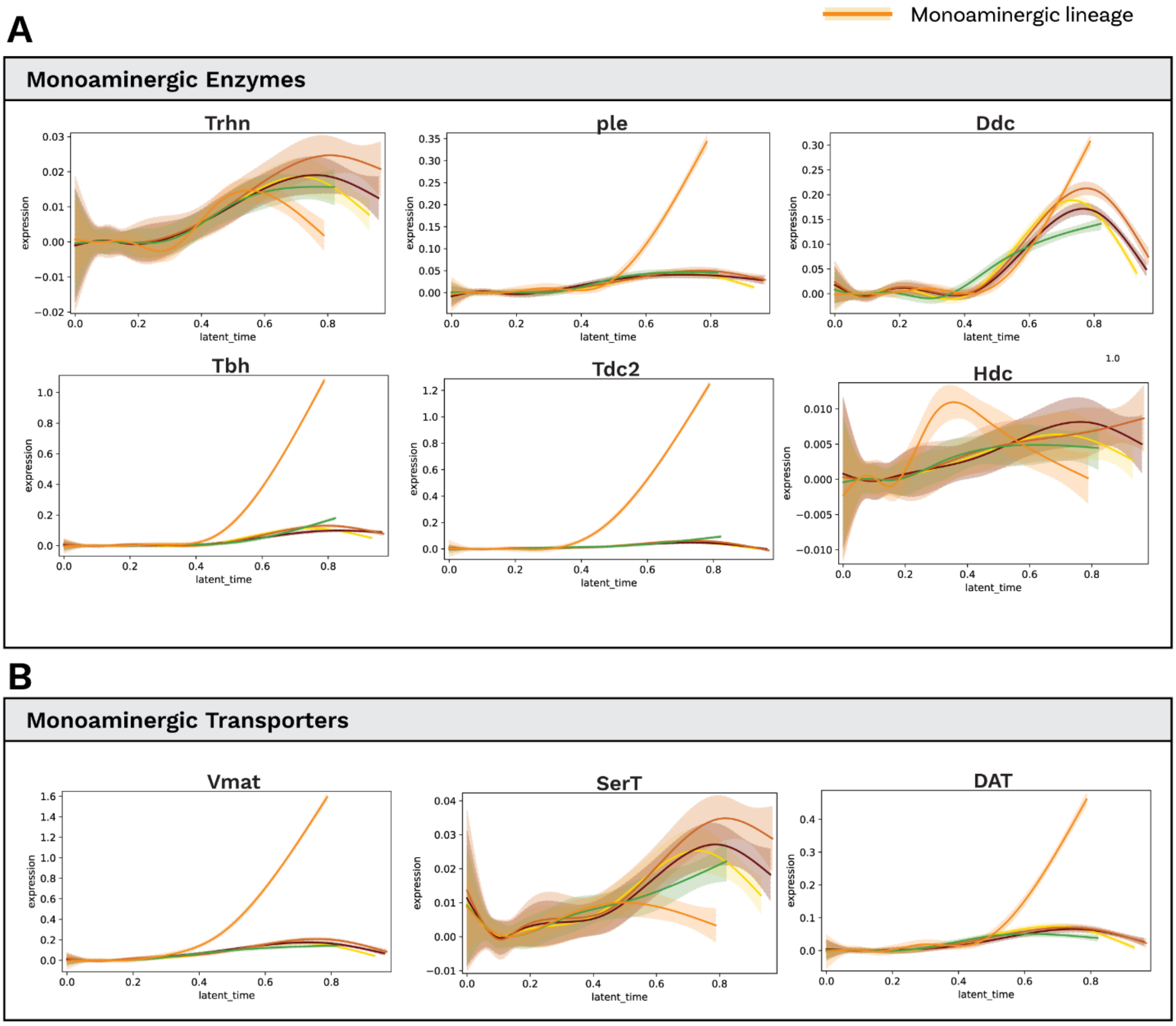


**Figure S6. Monoaminergic pathway gene trajectories along latent time in the neurogenic lineage**

**A. Monoaminergic enzymes.** Smoothed expression trends across latent time for major neuronal lineages (Acetylcholine, GABA_1/2, Glutamate, Monoamine). The monoaminergic lineage is highlighted in orange with a light-orange ribbon. Genes shown: *Trh*n, *ple*, *Ddc*, *Tbh*, *Tdc2*, *Hdc*. In the monoaminergic lineage, *ple* and *Ddc* (dopamine synthesis) and *Tdc2* and *Tbh* (tyramine/octopamine synthesis) rise strongly at late latent time, whereas *Trhn* and *Hdc* show modest or transient changes. Non-monoaminergic lineages remain low or show only brief increases. **B. Monoaminergic transporters.** Trajectories for *Vmat* (vesicular monoamine transporter), *DAT* (dopamine transporter) and *SerT* (serotonin transporter). All three increase toward late latent time in the monoaminergic lineage; *Vmat* and *DAT* are minimal or transient in other lineages, while *SerT* exhibits a more variable profile**.** Traces show lineage-wise smoothed means; ribbons indicate uncertainty. Axes: scaled expression vs. latent time.

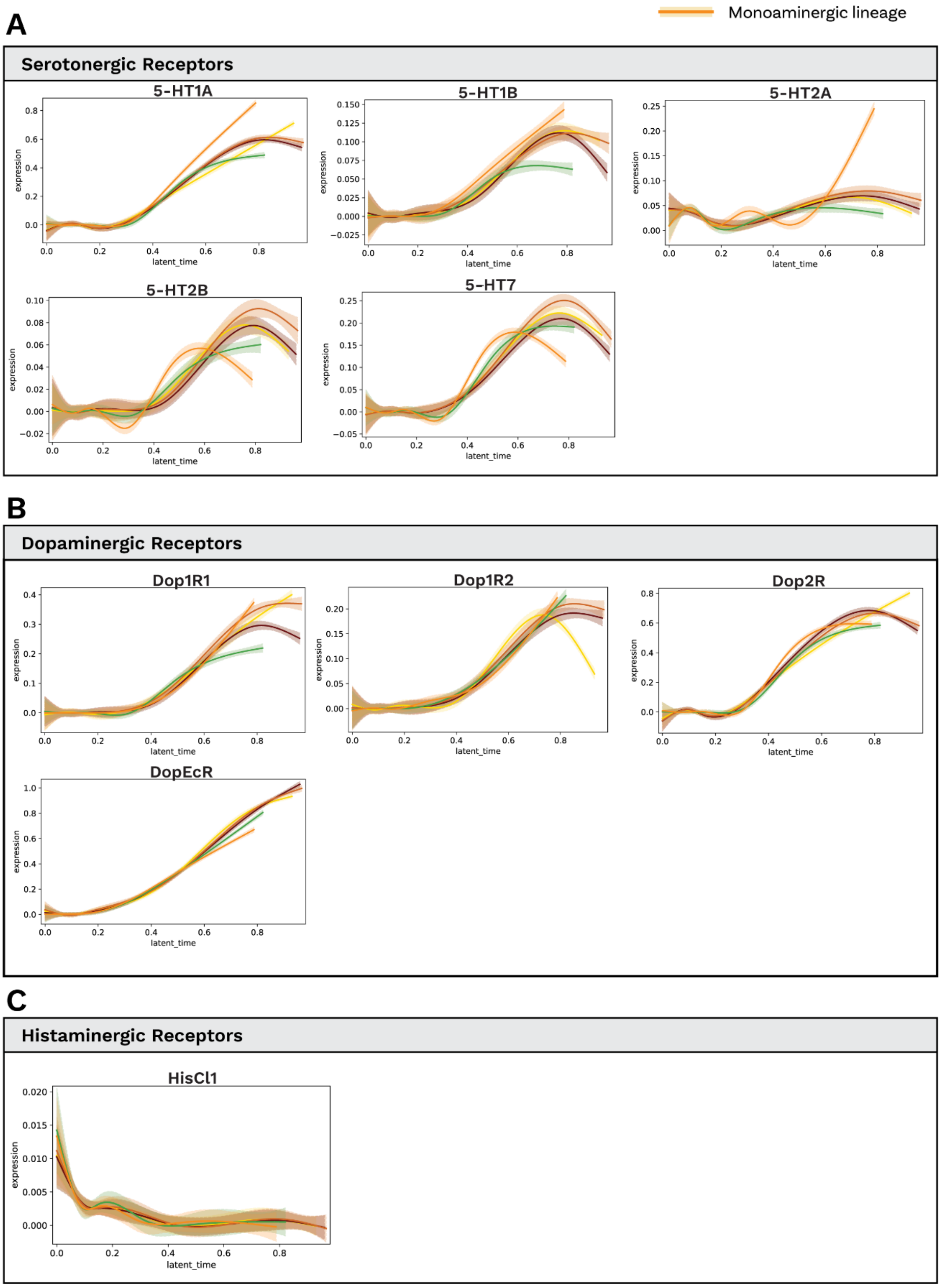
Monoaminergic lineage
A
Serotonergic Receptors
5-HT1A
5-HT1B
5-HT2A
5-HT2B
5-HT7
expression
latent_time
B
Dopaminergic Receptors
Dop1R1
Dop1R2
Dop2R
DopEcR
C
Histaminergic Receptors
HisCl1

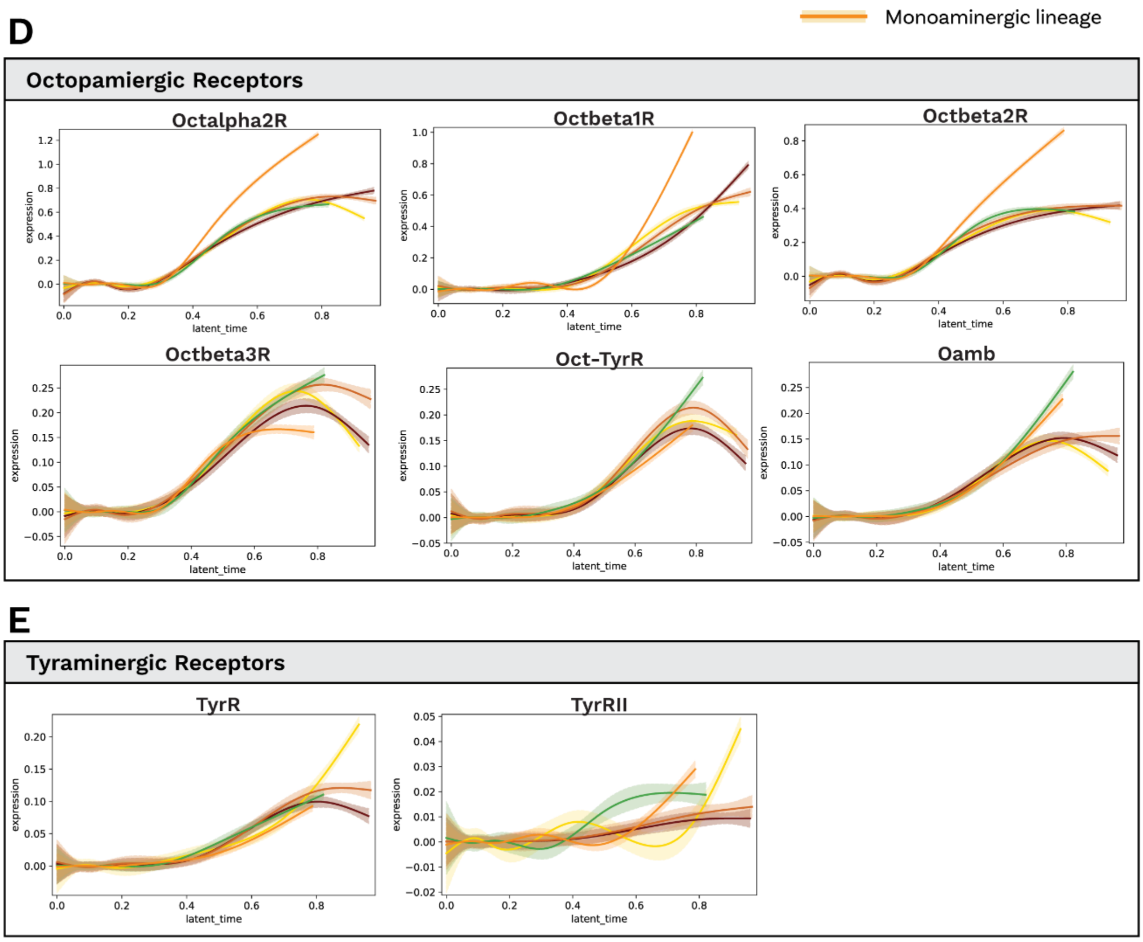


**Figure S7. Monoamine receptor trajectories across neuronal lineages and latent time**

Curves show lineage-wise smoothed expression across latent time for major neuronal lineages (Acetylcholine, GABA_1/2, Glutamate, Monoamine). The monoaminergic lineage is highlighted in orange with a light-orange ribbon indicating uncertainty. Axes: scaled expression (y) vs latent time (x).

**A. Serotonergic receptors. B. Dopaminergic receptors. C. Histaminergic receptors. D. Octopaminergic receptors. E. Tyraminergic receptors.**

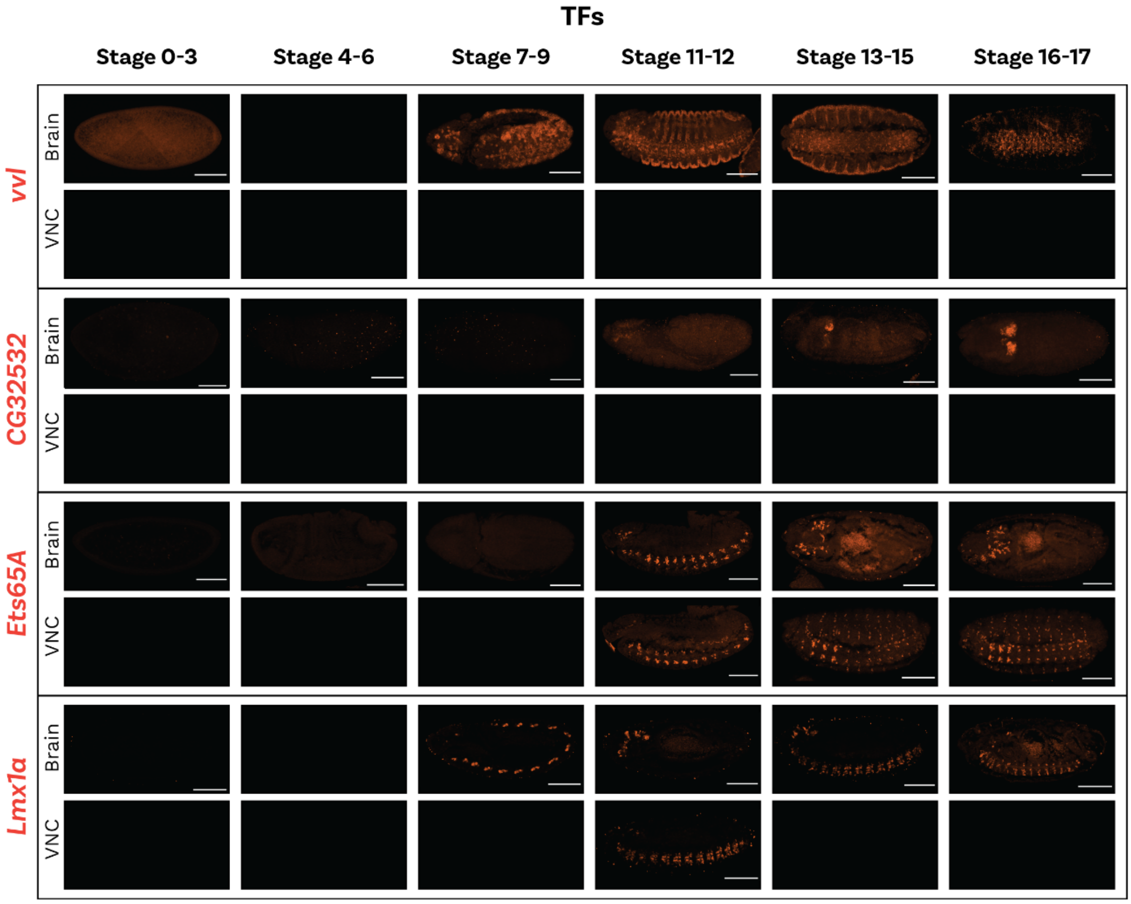


**Figure S8. HCR of candidate TFs across the monoaminergic lineage**

Whole-mount HCR staining across six embryonic windows (stages 0–3, 4–6, 7–9, 11–12, 13–15, 16–17). For each TF, the top row shows the brain and the bottom row the ventral nerve cord (VNC). Images are lateral views; anterior left. White scale bars shown (100 µm). *Vvl*, *CG32532*, *Ets65A*, *Lmx1a*. Signal is absent before gastrulation and then becomes detectable in patterned CNS domains from stage ~7–9 onward. By stages 11–12 and 13–15 each TF exhibits characteristic spatial enrichment—e.g., focal brain expression (*CG32532*), segmental VNC stripes (*Ets65A*), and combined brain/VNC domains (*Lmx1a*)—that persist until stage 16–17.

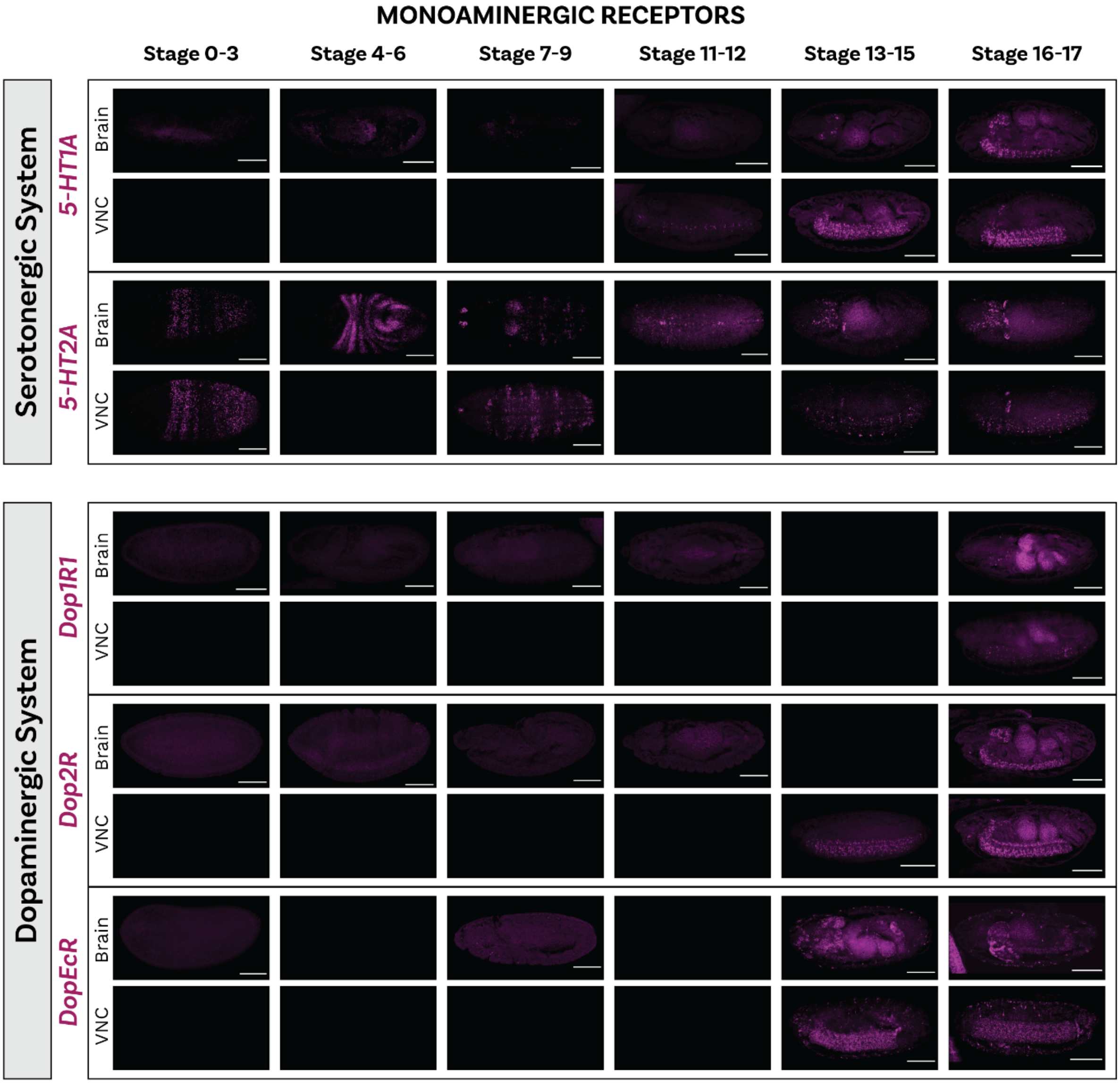


**Figure S9. HCR of serotonergic and dopaminergic receptors across embryogenesis**

Whole-mount HCR staining for serotonergic and dopaminergic receptors across six windows (stages 0–3, 4–6, 7–9, 11–12, 13–15, 16–17). For each gene, the top row shows the brain, the bottom row the ventral nerve cord (VNC). Images are lateral views; anterior left. Scale bars indicated (100 µm). **Serotonergic receptors.** *5-HT1A* is undetectable early and emerges in the CNS from stage 13-15, strengthening through stages 16–17 in brain and segmental VNC domains. *5-HT2A* shows early, widespread expression (from stages 0-3), including non-neural tissues, and later consolidates within CNS territories. **Dopaminergic receptors.** *Dop1R1* and *Dop2R* become detectable in the CNS from stages 16–17. *DopEcR* exhibits slightly broader CNS expression from stage 13-15 onwards.

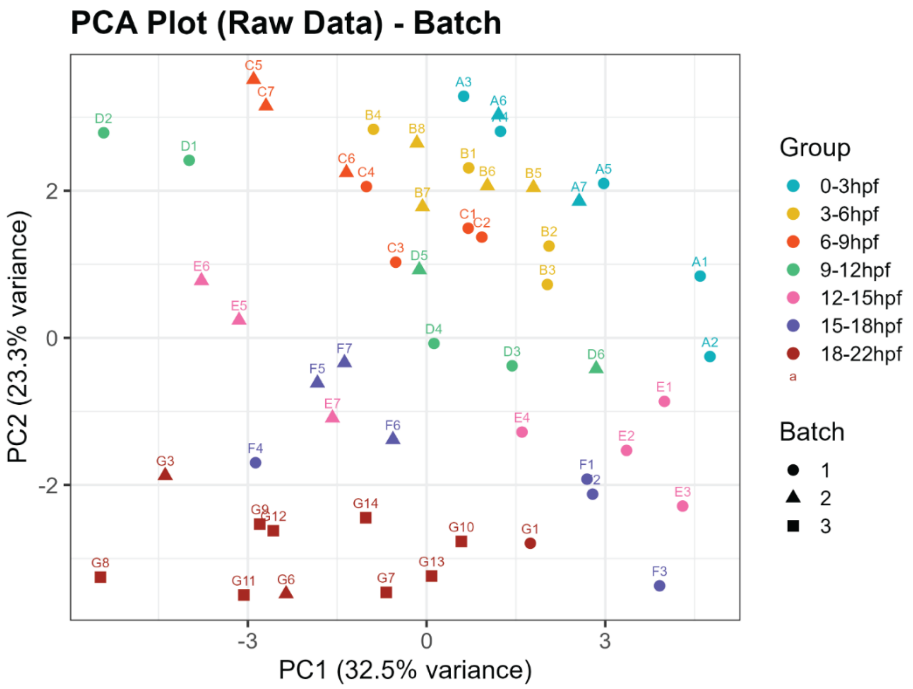


**Figure S10. PCA of targeted mass spectrometry data by developmental window and batch**

Principal components analysis of the raw measurements across seven time windows (0–3 hpf, 3–6 hpf, 6–9 hpf, 9–12 hpf, 12–15 hpf, 15–18 hpf, 18–22 hpf). Each point is a sample (labels denote sample IDs). Colour encodes developmental Group; shape encodes Batch (● batch 1, ▲ batch 2, ■ batch 3). Axes show the variance explained (PC1 = 32.5%, PC2 = 23.3%). Samples separate primarily by developmental window, with batches intermingled.

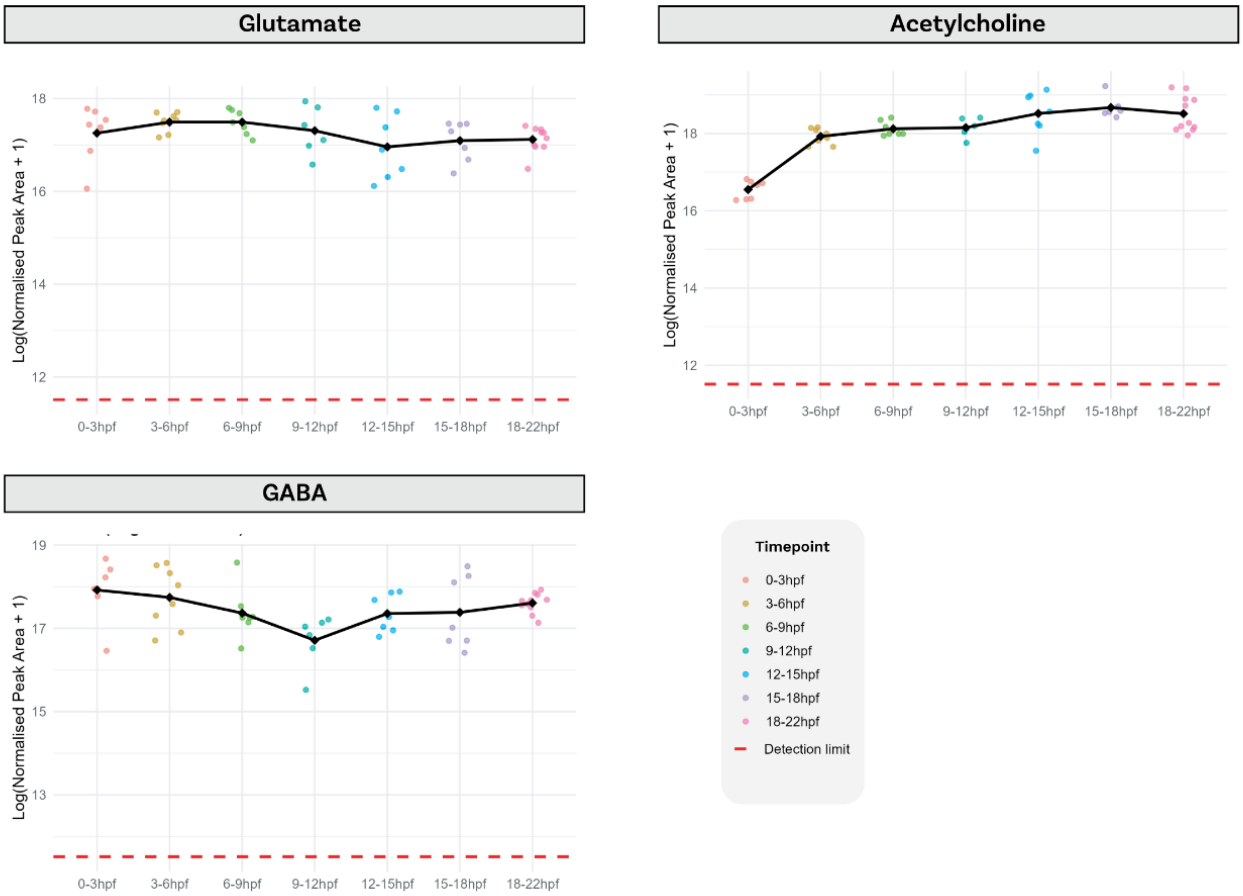


**Figure S11. Targeted mass spectrometry of other neurotransmitters**
Levels of glutamate, acetylcholine, and GABA were quantified across indicated timepoints using targeted mass spectrometry. Data are presented as mean from n ≥ 6 biological replicates.

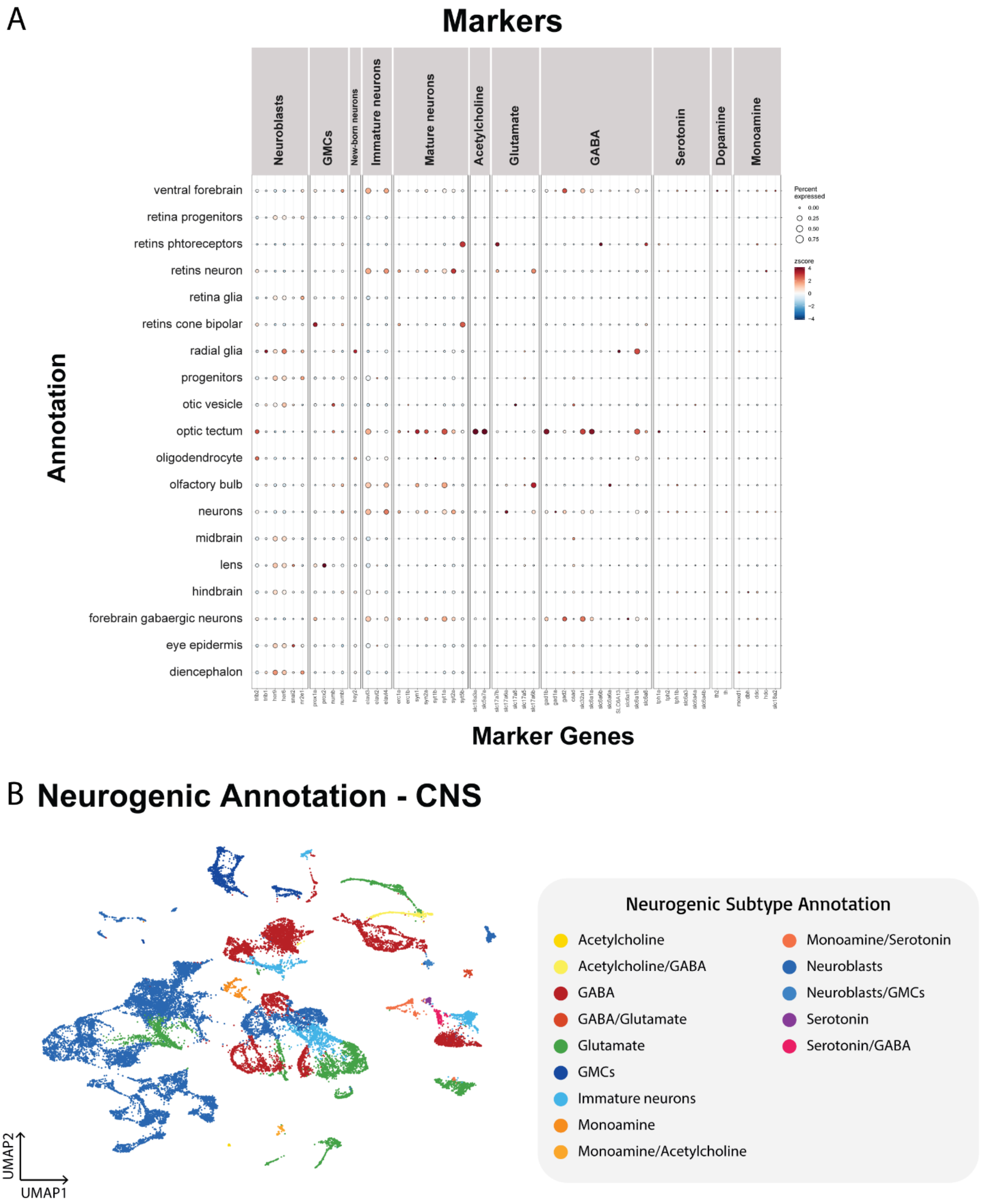


**Figure S12. Zebrafish CNS neurogenic landscape—marker concordance and subtype annotation**

**A. Marker concordance.** Dot plot of orthologous marker genes evaluated across CNS-derived clusters. Rows show anatomical/lineage annotations (e.g., diencephalon, forebrain GABAergic neurons, hindbrain, retina, neuroblasts), and columns group markers by neuronal state (Neuroblasts, GMCs, Immature/Mature neurons) and transmitter systems (Acetylcholine, Glutamate, GABA, Serotonin, Dopamine, Monoamine and mixed categories). Dot size indicates the fraction of cells expressing the gene in a cluster; colour is the z-scored average expression. **B. Neurogenic subtype annotation (CNS)**. UMAP of CNS cells coloured by inferred neurogenic/neuronal subtype. Labels denote transmitter classes and developmental states (Acetylcholine, Acetylcholine/GABA, GABA, GABA/Glutamate, Glutamate, Monoamine, Monoamine/Acetylcholine, Monoamine/Serotonin, Serotonin, Serotonin/GABA, Neuroblasts, Neuroblasts/GMCs, GMCs, Immature neurons).

**A**

**Monoamine**

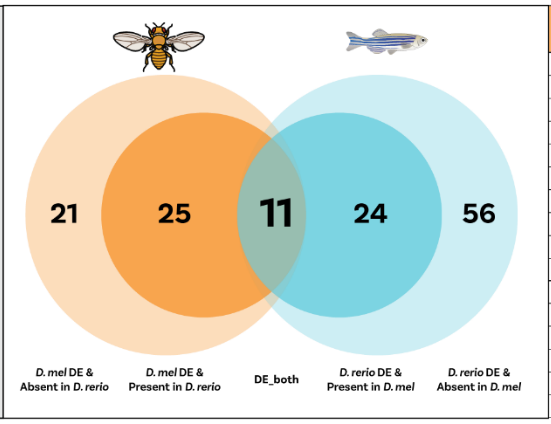


**Conserved TF Orthogroups**

| ***D. melanogaster*** | ***D. rerio*** |
|---|---|
| *cic* | *cica* |
| *Nfl* | *nfia; nfixb* |
| *cbt* | *klf11b; klf11a* |
| *CG4328; Lmx1a* | *lmx1bb; lmx1al; lmx1ba* |
| *FoxP* | *foxp4* |
| *trh* | *NPAS3* |
| *Eip78C* | *nr1d1* |
| *net* | *toh8* |
| *tj* | *mafa; mafb* |
| *Pdp1* | *hlfa* |
| *luna* | *klf6a* |

**B**

**Acetylcholine**

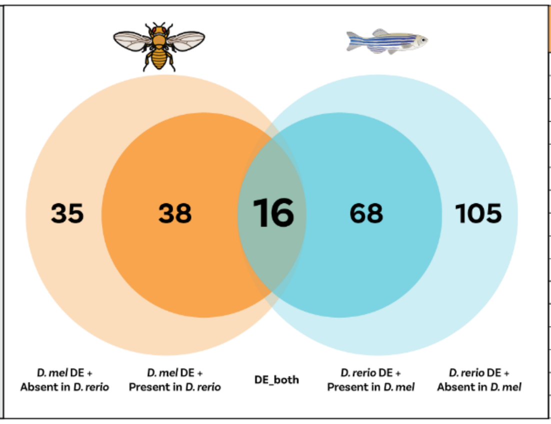


| ***D. melanogaster*** | ***D. rerio*** |
|---|---|
| *Mef2* | *mef2ca;mef2cb* |
| *Kr* | *bcl6aa;bcl6ab* |
| *Hr78* | *nr2c1* |
| *salr* | *sall3a* |
| *onecut* | *onecut* |
| *CG8301;CG12299;crol;CG18476* | *znfl2a;si:dkey-199m13.4;znf1156;si:ch73-299h12.6;zgc:174315;blf;zgc:165515* |
| *ftz-f1* | *nr5a2* |
| *scramb1* | *plscr3b* |
| *FoxP* | *foxp1a* |
| *exex* | *mnx1* |
| *otp* | *otpb;otpa* |
| *Hmx* | *hmx3a* |
| *kn* | *ebf2;ebf3b* |
| *grn* | *gata3;gata2a* |
| *acj6* | *pou4f1* |
| *unc-4* | *uncx* |

**C**

**GABA**

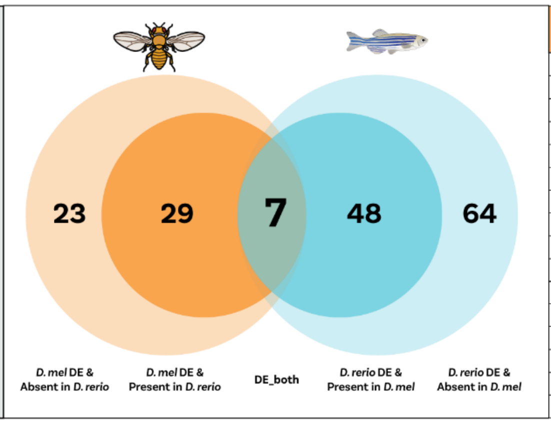


| ***D. melanogaster*** | ***D. rerio*** |
|---|---|
| *cic* | *cica* |
| *CG12605* | *scrt1a* |
| *Pur-alpha* | *puraa* |
| *CG9727* | *rfx7b* |
| *EcR* | *nr1h3* |
| *Pdp1* | *hlfa* |
| *Dll* | *dlx1a* |
| | *toh8* |

**D**

**Glutamate**

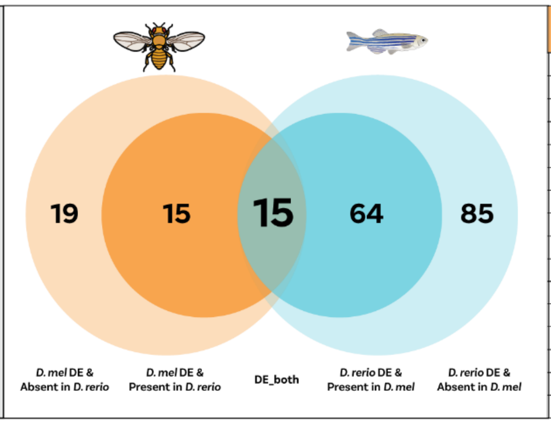


| ***D. melanogaster*** | ***D. rerio*** |
|---|---|
| *cic* | *cica* |
| *Mef2* | *mef2cb; mef2ab; mef2aa; mef2ca* |
| *tna* | *zmiz1a* |
| *scrt* | *scrt2;scrt1b;scrt1a* |
| *foxo* | *foxo3b;foxo3a* |
| *onecut* | *onecut1;onecut3b;onecut2;onecut3a* |
| *CG8301* | *zgc:101130;zgc:113348;BX470259.2* |
| *CG43689* | *myt1a;myt1la;myt1b* |
| *zfh1* | *zeb2b* |
| *Nfl* | *nfixb; nfia* |
| *CG13287* | *prdm8b* |
| *tsh* | *tshz2* |
| *Tusp* | *tulp4a* |
| *luna* | *klf7b;klf7a* |

**Figure S13. Cross-species conservation of transmitter-class TF programs**

**A–D.** EggNOG Orthogroup-level comparison of differentially expressed TFs between *D. melanogaster* (orange) and *D. rerio* (blue) within matched neurotransmitter classes: **A. Monoamine**, **B. Acetylcholine**, **C. GABA**, **D. Glutamate**. Venn diagrams summarise counts of Bilateria eggNOG orthogroups (OGs) containing TFs that are DE in the indicated class (Methods; adjusted $p < 0.05$, $|log2FC| \geq 0.25$). Numbers in each sector denote OGs unique to fly (left), unique to zebrafish (right), or shared (middle; DE in both species). The lower captions indicate whether the other species has a detectable member of the OG ("present") or lacks one in the proteome/annotations ("absent"). Tables to the right list representative conserved TF OGs (center intersection) with the corresponding gene members detected in each species; multiple names in a cell reflect many-to-one/one-to-many mappings within the same OG.
